# Microbiome assembly predictably shapes diversity across a range of disturbance frequencies

**DOI:** 10.1101/2021.08.02.454702

**Authors:** Ezequiel Santillan, Stefan Wuertz

**Author notes:** Correspondence to: Stefan Wuertz.

## Abstract

Diversity is frequently linked to the functional stability of ecological communities. However, its association with assembly mechanisms remains largely unknown, particularly under fluctuating disturbances. Here, we subjected complex bacterial communities in bioreactor microcosms to different frequencies of organic loading shocks, tracking temporal dynamics in their assembly, structure and function. Null modelling revealed a stronger role of stochasticity at intermediate disturbance frequencies, preceding the formation of a peak in α-diversity. Communities at extreme ends of the disturbance range had the lowest α-diversity and highest within-treatment similarity in terms of β-diversity, with stronger deterministic assembly. Stochasticity prevailed during the initial successional stages, coinciding with better specialized function (nitrogen removal). In contrast, general functions (carbon removal and microbial aggregate settleability) benefited from stronger deterministic processes. We showed that changes in assembly processes predictably precede changes in diversity under a gradient of disturbance frequencies, advancing our understanding of the mechanisms behind disturbance-diversity-function relationships.

## Introduction

Microbes exist typically as diverse, complex and dynamic communities^1^, responsible for all biogeochemical cycles worldwide^2^. These microbial communities or microbiomes provide crucial functions for global climate regulation, human health, biotechnology and bioremediation^3^. Microbial diversity is often related to community function^4^ and the ability to withstand environmental fluctuations that typically occur as disturbances^5^. Given the growing human population and its impact on natural and engineered ecosystems^6^, management and conservation practices are faced with increasing frequencies and magnitudes of various disturbances that occur on different scales. A concept of ecology that can be used to explore possible outcomes is the intermediate disturbance hypothesis (IDH), which predicts a diversity peak at intermediate levels of disturbance due to competition-colonization trade-offs faced by organisms^7^. Although the IDH has been influential in ecology^8^ and ecosystem conservation^9,10^, it is not a coexistence mechanism as initially thought^11^. Further, many studies have not found the diversity pattern predicted by the IDH^12,13^ and its relevance as a prediction tool is up for debate^14,15^. Therefore, studies are needed to address the mechanisms behind the observed disturbance-diversity relationships^16^.

Intermediate frequencies of exposure to a xenobiotic pollutant in our recent replicated sludge bioreactor study demonstrated higher α-diversity and relative influence of stochastic assembly compared to other exposure levels, after a short succession period of 35 days^17^. We hypothesized that when intermediate disturbance levels result in unpredictable environments where specialized traits are less advantageous to taxa, the stochastic equalization of competitive advantages would lead to a higher α-diversity, a causal relationship we named the intermediate stochasticity hypothesis (ISH)^17^. In contrast, either no disturbance or press disturbance conditions at the extreme ends of a disturbance range would allow fewer adapted organisms to dominate, thus lowering the α-diversity. Unlike the IDH, the ISH incorporates the role of assembly mechanisms as shapers of community structure (α- and β- diversity) across a disturbance gradient. Further, it predicts patterns not only in species richness but also in higher-order α-diversity indices, since variations in the underlying assembly mechanisms would affect abundance distributions of taxa. The ISH also considers that the output of a stochastic process is affected by some uncertainty, which means that there are several possible paths for the evolution of the structure and function of a community. In this regard, stochasticity operating at intermediate levels of disturbance in replicated systems could lead to similar high α-diversity (local, *e.g.*, within a reactor), but not necessarily to similar β-diversity (compositional variation across sites, *e.g.*, between reactors) and community function^17^. The idea of community assembly processes underlying the observed patterns of diversity is reasonable, as such processes are believed to shape community structure^18^, which also links them to ecosystem function. These processes, either deterministic or stochastic, are known to act in combination to form community assembly^19–22^ and can cause replicate communities to reach a similar or variable structure and function^17,23^. Further, while recent studies have reported positive correlations of strength of stochasticity with α-diversity in bacterial^24^ and fungal^25^ communities, the role of assembly processes behind diversity patterns under fluctuating disturbances is still unclear.

The objective of this work was to test the central tenet of the ISH that intermediate disturbance frequencies promote stochastic assembly processes, resulting in increased α-diversity and variable β-diversity^17^. We used an experimental system comprised of activated sludge sequencing batch reactors harboring complex microbial communities collected from a full-scale wastewater treatment plant. These were subjected to different frequencies of organic loading shocks, tracking temporal dynamics in their overall assembly, structure and function, without focusing on any particular taxa. The reactors had a working volume of 25 mL, representing a microcosm scale^26^. Replicates (n = 5) received double organic loading either never (L0, undisturbed), every 8, 6, 4 or 2 days (L1-4, intermediately-disturbed), or every day (L5, press-disturbed), for 42 days. Samples were analyzed using 16S rRNA gene metabarcoding and effluent chemical characterization. Patterns of α- and β-diversity were employed to assess temporal dynamics of bacterial community structure. Assembly mechanisms were quantified via null model analysis of phylogenetic turnover for each bioreactor.

## Results

### Intermediate disturbance frequencies exhibit higher taxonomic and phylogenetic α-diversity

Taxonomic α-diversity was evaluated using Hill diversity indices^27^ of orders zero (^0^D, taxa richness), one (^1^D) and two (^2^D), the latter being a robust estimate of microbial diversity^17^. Phylogenetic α-diversity was also considered through Faith’s phylogenetic distance^28^ unweighted (PD) and abundance-weighted (PD_W_). There was a temporal decrease in α-diversity for all disturbance frequency levels compared to the sludge inoculum for both taxonomic and phylogenetic α-diversity indices (Fig. 1A, Fig. S1). This drop was more pronounced, particularly within the first 14 days, when variability across same-level replicates was also highest. From d14 onwards, disturbance frequency had a significant effect on α-diversity (^2^D, Welch’s ANOVA P_adj_ = <0.001-0.015) (Fig. 1A). A peak in α-diversity at intermediate frequencies of disturbance was observed for all unweighted (^0^D, PD) and abundance-weighted (^1^D, ^2^D, PD_W_) indices evaluated in this study (Fig. 1A, Fig. S2, Fig. S3). Such parabolic pattern was significant from d21 onwards for ^2^D (Welch’s ANOVA P_adj_ ≤ 0.003), from d28 onwards for ^1^D (Welch’s ANOVA P_adj_ = 0.002-0.01), PD (Welch’s ANOVA P_adj_ = 0.003-0.037) and PD_W_ (Welch’s ANOVA P_adj_ = 0.005-0.013), and from d35 onwards for ^0^D (Welch’s ANOVA P_adj_ = 0.03-0.035).

**Fig. 1.**
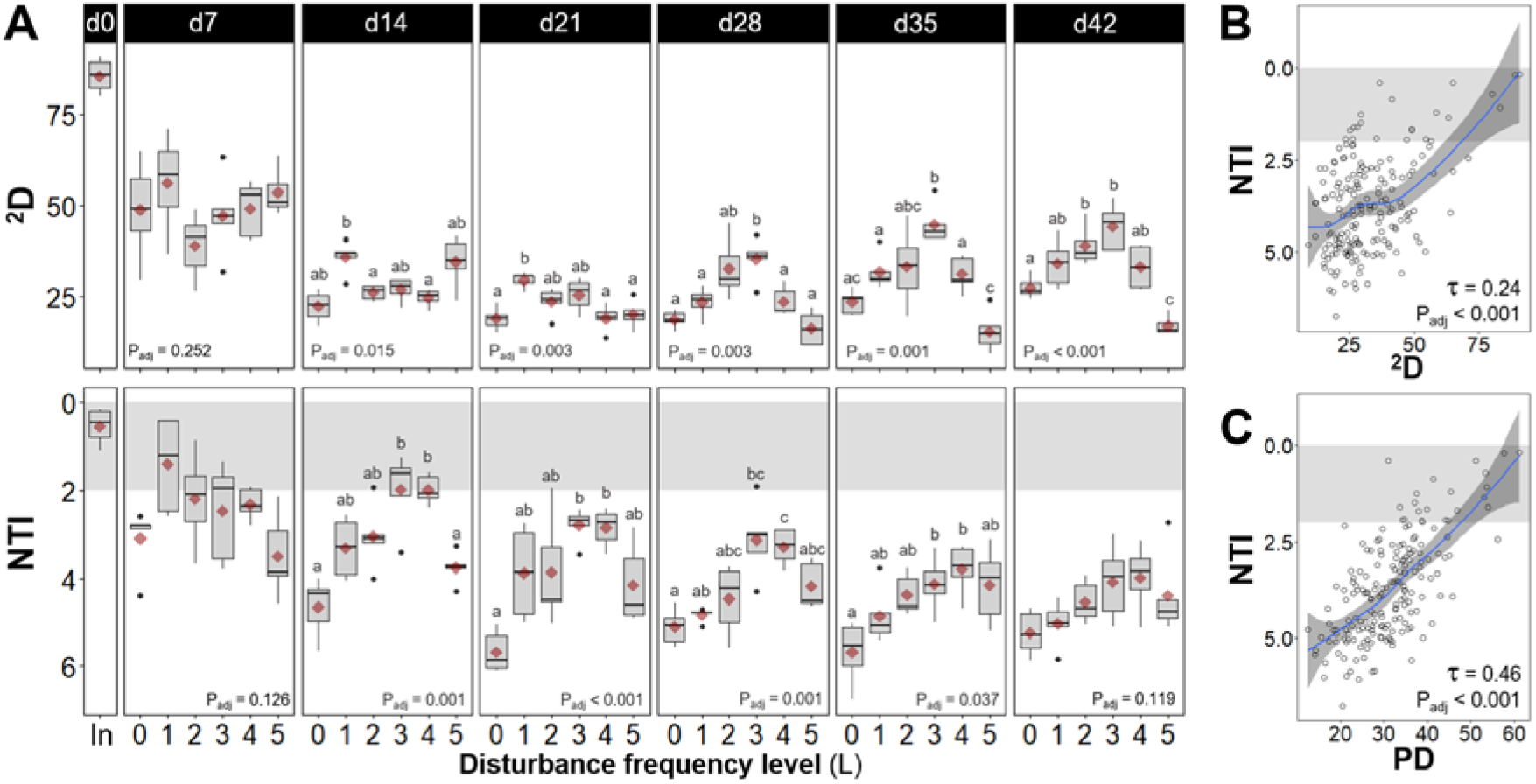
Community dynamics in α-diversity. (**A**) Community structure assessed via 2^nd^ order true α-diversity (^2^D, upper panels) and community assembly evaluated via the nearest taxon index (NTI, lower panels), from bacterial ASV data for different frequencies of organic loading disturbance (n = 5). Disturbance frequency levels (L): 0 (undisturbed), 1-4 (intermediately disturbed), 5 (press-disturbed). In: sludge inoculum (day 0, n = 4). Each panel represents a sampling day, red diamonds display mean values. Characters above boxes display Games-Howell post-hoc grouping (P_adj_ < 0.05). Welch’s ANOVA P-values adjusted at 5% FDR shown within panels. Correlations of (**B**) ^2^D and (**C**) phylogenetic diversity (PD) versus NTI from bacterial ASV data across all frequency levels and time points evaluated in this study (m = 184). Kendall correlation τ- and adjusted P-values are indicated within the panel. Blue line represents locally estimated scatterplot smoothing regression (loess) with confidence interval in dark-grey shading. Note the inverted y-axis for NTI, as values closer to zero indicate a higher relative contribution of stochastic assembly. Shaded in grey is the zone of significant stochastic phylogenetic dispersion, |NTI| < 2.

### Community assembly temporal dynamics precede α-diversity patterns across disturbance frequencies

Assembly processes were first evaluated by modelling the phylogenetic dispersion of a given community against the null expectation, through the nearest taxon index (NTI)^29^. We observed higher stochasticity at the initial stages of the experiment (d0-14), which decreased in relative intensity over time across disturbance levels for both unweighted (NTI) and abundance-weighted (NTI_W_) indices (Fig. 1A, Fig. S2). There was a stronger role of stochastic assembly processes at intermediate disturbance frequencies as shown by NTI values closer to zero (*i.e.*, lower |NTI| values); this was significant from d14 onward (NTI Welch’s ANOVA P_adj_ = <0.001-0.037) but was reduced towards the end of the study becoming non-significant on d42. Games-Howell post-hoc grouping indicated that the parabolic pattern of NTI across disturbance frequency levels preceded (d14-35) the formation of a peak in α-diversity (d21-42) at intermediate levels of disturbance, with two to three groups significantly differentiated (Fig. 1A). Stochastic assembly processes were less prevalent when abundance weighing was included in the calculation of the NTI index (NTI_W_), meaning that phylogenetic dispersion compared to the null expectation was higher among individual organisms than it was among taxa. Nonetheless, there was a significant peak in NTI_W_ values at intermediate frequencies of disturbance on d7 and d14 (NTI_W_ Welch’s ANOVA P_adj_ = 0.001). This parabolic pattern of NTI_W_ was evident on d7, preceding that of NTI, but disappeared on d21 and reverted from d28 onwards. Also, significant phylogenetic signals were observed via mantel correlogram analysis (Fig. S5) mostly across relatively short phylogenetic distances, justifying the use of phylogenetic null modelling to evaluate community assembly processes in this study.

Stochastic assembly was more important when α-diversity was higher, particularly for phylogenetic diversity. This was shown by significant Kendall correlation τ values (0.24-0.46, P_adj_ < 0.001) between NTI and α-diversity indices (Figs. 1B-C, Fig. S4). Kendall correlation τ values were also positive (0.20-0.26) and significant (P_adj_ < 0.001) between NTI_W_ and phylogenetic α-diversity indices (PD, PD_W_) and unweighted taxonomic α-diversity (^0^D), but not between NTI_W_ and abundance-weighted taxonomic α-diversity (^1^D, ^2^D) (Fig. S4). The estimation of all the aforementioned indices over time using rarefied ASV sequencing data yielded the same significant patterns via Welch’s ANOVA, with the exception of NTI_W_ on d21 and d42 (see supplementary file).

### β-diversity patterns display similarity at low and high disturbance frequencies and higher variability at intermediate ones

Community structure in terms of β-diversity showed temporal changes, which varied across disturbance levels for both Unifrac phylogenetic distances (Fig. 2A) and Bray-Curtis taxonomic distances (Fig. 2B). Unconstrained ordination displayed a dispersion effect in overall community structure over time, particularly after 7 days, with communities in each reactor diverting from each other (Fig. 2A). To disentangle the effect of disturbance from temporal dynamics, constrained ordination via canonical analysis of principal coordinates (CAP) was used at each time point (Fig. 2B). Group-average cluster similarity (60%) was included to detect formations of clusters of community structure. Differences in β-diversity across disturbance levels were statistically significant at all time points evaluated (PERMANOVA P_adj_ < 0.001), without significant effects of heteroscedasticity (PERMDISP P_adj_ > 0.14) (Table S1). Replicate reactors at the undisturbed (L0) and press-disturbed level (L5) clustered separately from intermediate disturbance levels on all sampling days (except on d7 and d21 for L0) (Fig. 2B), both levels having 0% misclassification error at all time points assessed (Fig. 2C). Comparatively, reactors at intermediate disturbance frequencies (L1-4) clustered together and showed higher dispersion across replicates within the same level, with CAP misclassification errors above zero (Fig. 2B-C). Thus, replicate reactors were less similar to each other at intermediate levels of disturbance, while replicates at low (undisturbed) and high (press-disturbed) disturbance frequencies were more similar. Likewise, community assembly assessed via the beta nearest taxon index (βNTI)^30^ showed a higher relative contribution of stochasticity at intermediate levels of disturbance (Fig. 2D), with βNTI values closer to zero, indicating that phylogenetic turnover across within-treatment replicates was closer to the null expectation. Similarly to what we observed through the NTI, the relative importance of stochasticity decreased with time in the experiment (*i.e.*, higher |βNTI| values) and when abundance weighing was included in the calculation of the βNTI values (βNTI_W_) (Fig. S6). The observed temporal changes in bacterial community structure across disturbance frequencies were consistent with phylum-level dynamics in relative abundances (Fig. S7), although the focus of this study was on overall community dynamics and not on any particular group of taxa.

**Fig. 2.**
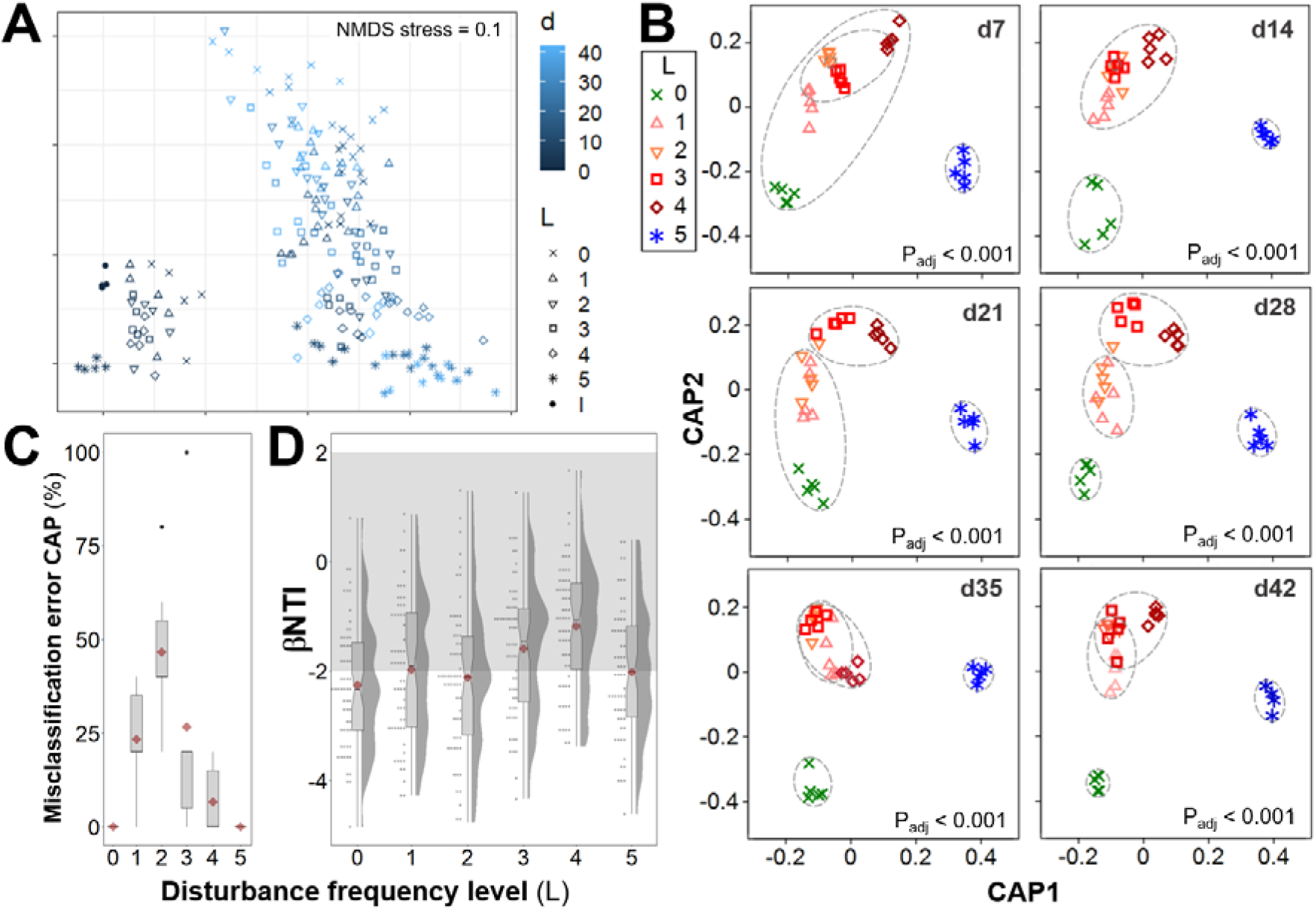
Temporal dynamics of β-diversity community structure and assembly for bacterial ASV data across different frequencies of organic loading disturbance (n = 5 bioreactors). (**A**) Unconstrained NMDS ordination (weighed Unifrac β-diversity, Hellinger transformed data) for all 184 samples collected. Disturbance frequency levels (L): 0 (undisturbed), 1-4 (intermediately disturbed), 5 (press-disturbed). I: Sludge inoculum (day 0, n = 4). (**B**) Constrained canonical analysis of principal coordinates (CAP) ordinations (Bray-Curtis β-diversity, squared root transformed data) on different sampling days, including ellipses of 60% group-average cluster similarity and PERMANOVA adjusted P-values. (**C**) Misclassification errors at each disturbance frequency level, via the leave-one-out allocation of observations to groups from CAP at each time point after d0 (n = 6 sampling days). Bray-Curtis β-diversity, squared root transformed data. Red diamonds display mean values. (**D**) Beta nearest taxon index (βNTI) at each disturbance frequency level, from pairwise comparisons across within-treatment replicates at each time point after d0 (n = 60 comparisons). Red diamonds display mean values. Notches show the 95% confidence interval for the median. When notches do not overlap, the medians can be judged to differ significantly. Shaded in grey is the zone where stochastic processes significantly dominate, |βNTI| < 2. βNTI values closer to zero indicate a higher relative contribution of stochastic assembly.

### Community function dynamics and correlations with community structure and assembly

Bacterial community function was assessed over time via influent chemical oxygen demand (COD) removal, sludge volume index (SVI), and influent total Kjeldahl nitrogen (TKN) removal, as measure of carbon removal, sludge settleability and nitrogen removal, respectively (Fig. 3A). Carbon removal and sludge settleability, which are functions associated with a broad range of taxa (*i.e.*, general functions), improved over time during the experiment. High carbon removal (> 0.97) was achieved at all disturbance frequency levels from d21 onwards, with no significant differences on days 35 and 42, after a period of high variability for same-level replicates during the first 14 days. Sludge settleability increased with disturbance frequency, with undisturbed (L0) reactors showing the lowest settleability from d21 onwards and intermediately disturbed levels reaching the highest settleability on d42 (SVI Welch’s ANOVA P_adj_ = 0.018). The nitrogen removal function (TKN removal), which is related to specialized bacteria (ammonia oxidizers), significantly differed across disturbance frequencies (TKN removal Welch’s ANOVA P_adj_ < 0.001) with the highest removal at intermediately disturbed levels during the first 21 days. From d28 onwards, L0 to L4 reactors had similarly high average nitrogen removal (> 0.9), and only the press disturbed reactors (L5) continued to have lower nitrogen removal (< 0.7) than that of the initial sludge inoculum (0.8). Effluent values of TKN, ammonia, nitrite and nitrate showed that TKN removal occurred via nitrification (Fig. S8).

**Fig. 3.**
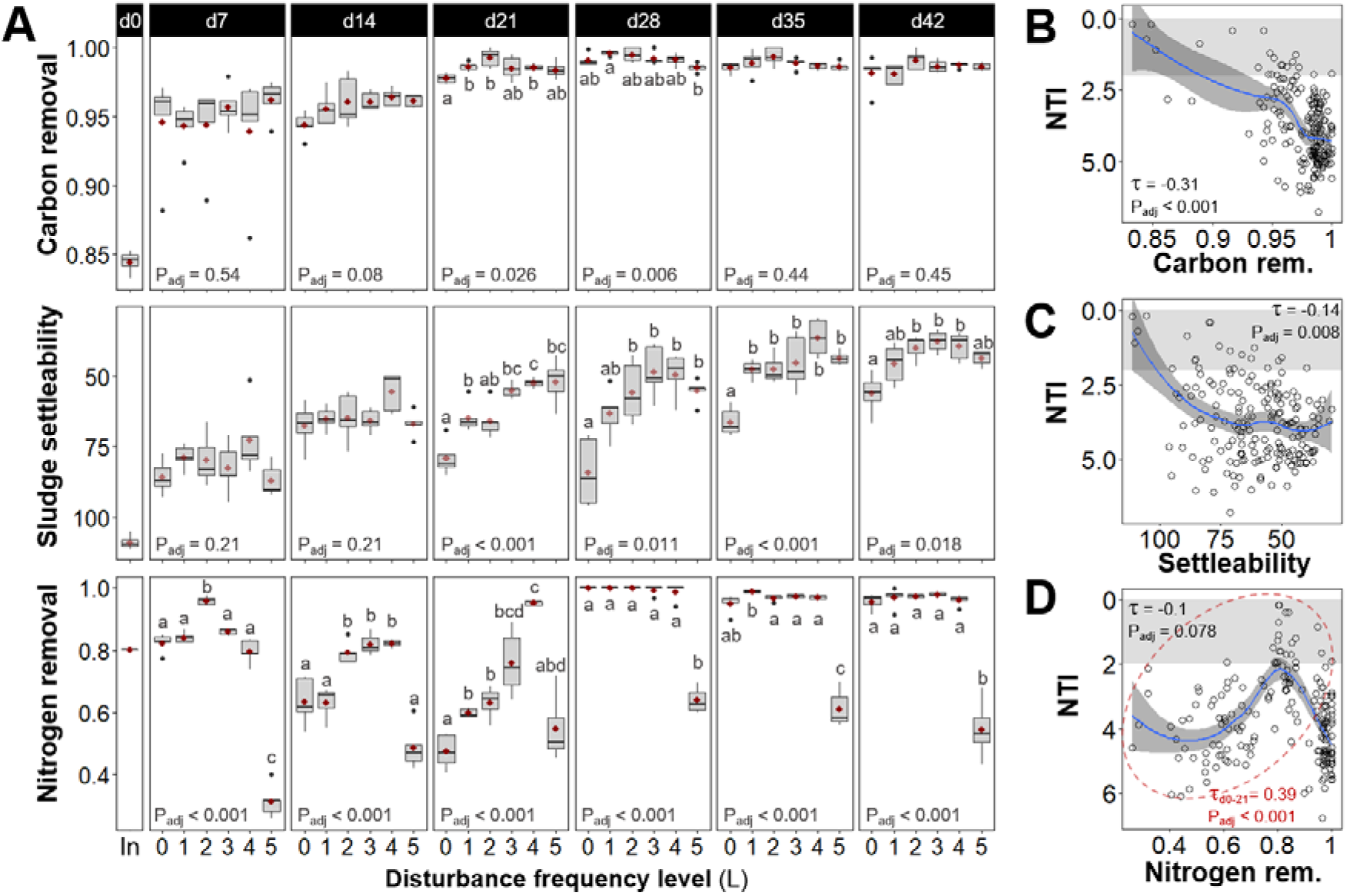
Community function assessed via influent chemical oxygen demand removal (carbon removal, upper panels), sludge volume index (sludge settleability, middle panels), and influent total Kjeldahl nitrogen removal (nitrogen removal, lower panels) for different frequencies of organic loading disturbance (n = 5). Disturbance frequency levels (L): 0 (undisturbed), 1-4 (intermediately disturbed), 5 (press-disturbed). In: sludge inoculum (day 0, n = 4). Each panel represents a sampling day, red diamonds display mean values. Characters above boxes display Games-Howell post-hoc grouping (P_adj_ < 0.05). Welch’s ANOVA P-values adjusted at 5% FDR shown within panels. Correlations of (**B**) carbon removal, (**C**) sludge settleability, and (**D**) nitrogen removal, versus NTI from bacterial ASV data across all frequency levels and time points evaluated in this study (m = 184). Kendall correlation τ- and adjusted P-values are indicated within the panels. Blue line represents locally estimated scatterplot smoothing regression (loess) with confidence interval in dark-grey shading. Shaded in grey is the zone of significant stochastic phylogenetic dispersion, |NTI| < 2. Red ellipse and τ- and P-value in panel (**D**) indicate data at initial stages of succession (d0 to d21). Note the inverted axis for sludge settleability, as it increases with decreasing SVI values, and for NTI, since values closer to zero indicate a higher relative contribution of stochastic assembly.

Carbon removal had an overall significant negative Kendall correlation with α-diversity indices (τ < −0.21, P_adj_ < 0.001), whereas sludge settleability and nitrogen removal showed non-significant correlations with α-diversity across the study (Fig. S9). Correlations between general functions of carbon removal and sludge settleability and both NTI and NTI_W_ were negative and significant across all time points and disturbance frequencies of the study (Fig. 3B-C, Fig. S9), indicating improved performance of these functions under stronger deterministic assembly mechanisms. Nitrogen removal had a non-significant overall correlation with NTI and NTI_W_ (Fig. 3D, Fig. S9), which became positive and significant when only the first 21 days of the study were considered (NTI τ_d0-21_ = 0.39, P_adj_ < 0.001, Fig. 3D; NTI_W_ τ_d0-21_ = 0.46, P_adj_ < 0.001, Fig. S10), suggesting better performance of this function under higher stochastic assembly throughout the initial weeks of the study.

## Discussion

In this study we found stochastic assembly processes to be more important at intermediate disturbance frequencies where the highest α-diversity was also observed, together with high β- diversity dispersion across within-treatment replicates as predicted by the ISH^17^. Furthermore, we showed that a peak in the relative contribution of stochasticity preceded a peak in α-diversity across a disturbance frequency range. Also, we observed that higher stochasticity during initial successional stages correlated with better nitrogen removal (specialized function) at intermediate disturbance frequencies, while carbon removal and microbial aggregate settleability (general functions) improved in step with more deterministic forces. These findings highlight the utility of the ISH for a mechanistic understanding of disturbance-diversity-function relationships.

We expanded our earlier work^17^ by using a different type of disturbance and microbial community inoculum, a relevant scenario given the multidimensional nature of disturbance^31^. Employing taxonomic and phylogenetic diversity metrics, in both unweighted and abundance-weighted forms, allowed us to cover a broader aspect of α-diversity. Taxonomic resolution was also improved by the use of amplicon sequence variants (ASVs) compared to operational taxonomic unit (OTU) clustering^32^ with about one to two orders of magnitude fewer spurious units^33^, allowing for a better estimation of unweighted α-diversity (*i.e.*, taxa richness). We further verified that the observed patterns occurred independently of data rarefaction, given the lack of consensus about this practice^34^ and the fact that it is known to affect (mainly unweighted) estimations of α-diversity^35^. Assembly processes were tracked over time using a phylogenetic null modelling methodology, which has been tested and recommended in microbial ecology^20,30,36^. Additionally, general and specific functions were evaluated against structure and assembly. All the aforementioned enhancements allowed us to test the ISH, while also gaining new insights into the role of assembly processes behind disturbance-induced changes in community structure and function over time.

Our experimental system produced a succession scenario in which bacterial communities had to adapt to change from a full-scale system to a bioreactor microcosm setup along a disturbance frequency gradient, similarly to what we described in a prior study^37^. With regards to community structure, succession led to a significant hump-backed pattern of α-diversity for all composition- and abundance-based indices employed in the study, which occurred after 21 days for ^2^D, 28 days for ^1^D, PD and PD_W_, and 35 days for ^0^D. Thus, the observed dynamics in community structure were stronger in terms of relative abundances than richness (^2^D, ^1^D vs. ^0^D), as well as at the phylogenetic versus taxonomic level (PD vs. ^0^D). The appearance of higher phylogenetic α-diversity at intermediate levels of disturbance for both unweighted (PD) and abundance-weighed (PD_W_) indices suggests that considering evolutionary relationships among organisms^38^ could also aid in assessing the effect of varying disturbances on community structure under succession. In our study, disturbance promoted the co-occurrence of phylogenetically distinct organisms, suggesting that additional niches were created at intermediate disturbance frequencies that were occupied by ecologically different species, thus reducing competitive exclusion. Conversely, phylogenetic clustering at undisturbed and press-disturbed levels can be interpreted as communities structured by environmental filtering^29^. Additionally, temporal analysis of community structure in terms of β-diversity revealed three different clusters for undisturbed, press-disturbed and intermediately disturbed reactors. Further comparison of replicates within the same disturbance frequency level showed higher β-diversity variability at intermediate disturbance levels, which was coherent with prior observations in freshwater ponds^39^ and sludge bioreactors^17^ where β-diversity increased with stochastic assembly. Our findings are relevant for understanding disturbance-diversity relationships, since few studies have reported parabolic α-diversity patterns using abundance-based indices^8^. Furthermore, variations in β-diversity among ecological communities that are subject to large and fluctuating disturbances are believed to provide insights about the mechanisms driving changes in α-diversity and function^40^.

We observed similar trends of phylogenetic dispersion within a single community (NTI) and the phylogenetic turnover between communities of the same treatment level (βNTI), compared to the null expectation. Stochasticity was more important during initial successional stages of the study, with initial NTI and βNTI values closer to zero (*i.e.*, closer to the null expectation of the model). Relatively, the overall strength of deterministic processes increased with time, with higher |NTI| and |βNTI| values. Similarly, late succession stages were shown to be governed by deterministic processes in plant forest^41^ and microbial groundwater communities^42^. Furthermore, α-diversity-based temporal assembly dynamics revealed a parabolic pattern in NTI and NTI_W_, through the disturbance frequency gradient, which was evident after 14 and 7 days of the study, respectively, before the appearance of similar parabolic patterns across various α-diversity indices. This preceding pattern is considered here as a strong indicator of assembly mechanisms operating to shape community structure. It is, therefore, plausible that stochastic assembly mechanisms were first favored at intermediate disturbance frequencies, prompting subsequent changes of community structure that resulted in the observed higher α-diversity as the ISH proposes^17^. These observations are also coherent with the idea that secondary succession is community assembly in action^43^. The disturbance range in this study produced different secondary succession scenarios, with communities in the sludge of each bioreactor likely experiencing different re-colonization processes from their bacterial seed-bank (*i.e*, low-abundance or rare taxa), via stochastic processes such as priority effects^44^ followed by historical contingency^45^ and legacy effects^3^. Importantly, external dispersal processes^46^ (*i.e.*, bacterial immigration) could not influence community assembly since bioreactors within this study were operated as closed systems. Indeed, microbial seed-banks are thought to contribute to the maintenance of microbial diversity^47^ and have been described as essential for understanding temporal community changes^48^. Further, stochastic assembly processes were shown to be more preponderant within the rare fraction of the microbial community^22^. Nonetheless, other processes might also be promoting stochastic assembly at intermediate disturbance frequencies, like ecological drift^36^ and feedback mechanisms linked to density dependence and species interactions^49^. Hence, a disturbance frequency gradient can not only result in nonlinearities for growth rates that would affect the outcome of competition^14,31^, it could also alter the relative contribution of stochastic and deterministic mechanisms of community assembly that underlie changes in community structure^17^. Furthermore, our results showed that, over a range of disturbance frequencies, assessing temporal community assembly patterns during succession can act as a sentinel of upcoming patterns of diversity.

Stochasticity was positively correlated with better nitrogen (as TKN) removal via nitrification at intermediate disturbance frequencies during the initial successional stages where stochastic processes were also generally prevalent. Nitrification functions are carried out by specific taxa (*i.e.*, nitrifiers), which are slow growers, nutritionally inflexible, sensitive to inhibitors and less phylogenetically diverse than many other key functional guilds^50^. Yet, the recruitment of nitrifying organisms from the microbial seed-bank was important for the recovery of nitrification, following the removal of a long-term disturbance of altering food-to-biomass and carbon-to-nitrogen ratios in sludge bioreactors, although resilience varied across identically treated replicates^51^. Also, partial recovery of nitrification in sludge bioreactors was observed at intermediate frequencies of 3-chloroaniline disturbance, where stochastic assembly processes and within-treatment variability were also higher^17^. Conversely, general functions of carbon removal and settleability performed better when deterministic processes were stronger (higher |NTI| values). Carbon removal was better when α-diversity was lower, similarly to what was reported previously using a different xenobiotic disturbance in bioreactors^17^. Hence, a more diverse community does not necessarily translate into better ecosystem functions^17,52^. Our data suggest that general functions thrive during stronger deterministic processes, while specialized functions might be favored by stochasticity at initial successional stages. Future studies assessing the effect of fluctuating disturbances on community diversity and function should also consider the type of function (*e.g.*, specific or general), the stage of succession after the disturbance, and the underlying assembly mechanisms.

The observed patterns in community assembly, structure and function were time-dependent. The ISH successional pattern appears to be transient, as assembly mechanisms across disturbance frequency levels were not significantly different towards the end of the study on d42, while α-diversity continued to display a significant parabolic pattern. If the gradient of disturbance frequencies is maintained over time, then the peak in α-diversity at intermediate levels might continue during the late successional stages, but this remains to be investigated. Nonetheless, most relevant bacteria in activated sludge have generation times of less than 24 h. Hence, the 42-day length of this study represented around tens to hundredths of generations of many different taxa, allowing the detection of significant patterns in assembly and structure. Further research in a variety of ecosystems is needed to validate the broad applicability of the ISH, particularly considering that disturbance can vary in type, frequency, intensity, driver and impact^31,53^. Studies at different scales are also necessary since ecological patterns can vary across spatial, temporal and phylogenetic scales^3^, while the effect of dispersal processes could also be evaluated within open systems.

Although a similar study on communities of larger organisms would require considerably larger scales of space and time, some modelling approaches suggest that ISH-like patterns (Fig. 4) could emerge in community assembly and structure under varying disturbances. For example, forest fire modelling showed that intermediate lightning strike frequency values yielded higher diversity with a close balance between stochastic and deterministic forces, which were highly sensitive to probabilistic events leading the system to diverse trajectories^54^. Likewise, a conceptual model developed for plants and animals suggested that high variation in resource abundance and location in space and time, which could be caused by disturbance, would favor diversity via adaptation through novelty and innovation (*i.e.*, stochasticity) generation^55^. The predictions of the ISH could help to identify cases when disturbance-induced stochastic assembly promotes alternative states of community structure that compromise or enhance ecosystem function, so as to design mitigation or intensification strategies. Furthermore, it could be used to promote community resistance and resilience to future disturbances via increased α-diversity and functional-gene diversity. Alternatively, this theoretical framework could help develop functionally resilient communities that do not occur naturally, through the stochastic mechanisms that are initially elicited at intermediate frequencies of disturbance. Therefore, we propose that the ISH has potential for a general understanding of disturbance-induced changes in community structure and function during succession, by integrating the influence of the underlying assembly processes over time.

**Fig. 4.**
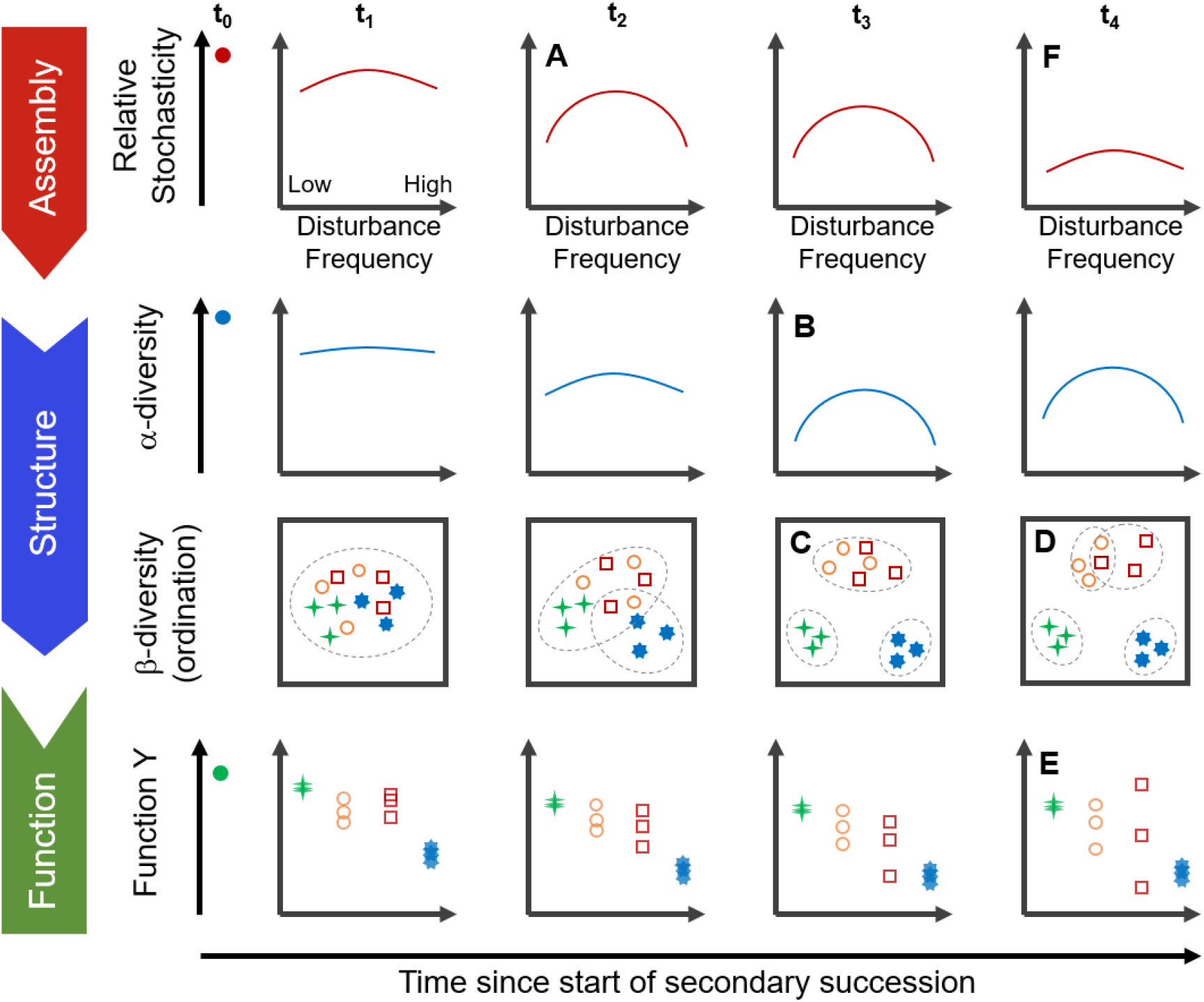
Conceptual representation of the intermediate stochasticity hypothesis (ISH) to describe patterns of assembly and structure along a disturbance frequency gradient, for communities in secondary succession (starting at time point t_0_). (**A**) Initially, stochastic assembly mechanisms (*e.g*., priority effects, historical contingency and legacy effects) are favored at intermediate disturbance frequencies, promoting re-colonization processes from the low-abundance fraction of the community or seed-bank. (**B**) Subsequently, these are followed by changes in the community structure that manifest as a peak of α-diversity at intermediate levels of disturbance. (**C**) At least three separated clusters of β-diversity ordination would form over time across the disturbance range. However, stochasticity operating at intermediate disturbance levels may lead to variable within treatment (**D**) β-diversity and (**E**) community function. (**F**) The overall relative contribution of stochasticity decreases with succession time. The observed patterns of diversity are stronger in terms of relative abundances than richness, as well as at the phylogenetic versus the taxonomic level.

## Materials and Methods

### Experimental design and function analyses

We employed 30 sequencing batch bioreactors at a microcosm scale (25-mL working volume), inoculated with activated sludge from a full-scale wastewater treatment plant in Singapore and operated for 42 days at 30°C in an incubator shaker. The daily complex synthetic feeding regime (adapted from Santillan *et al.*^51^) included double organic loading at varying disturbance frequencies. Six levels of disturbance were set in quintuplicate independent reactors (n = 5), which received double organic loading either never (undisturbed), every eight, six, four, or two days (intermediately-disturbed), or every day (press-disturbed). Level numbers were assigned from 0 to 5 (0 for no disturbance, 1 to 5 for low to high disturbance frequency). Disturbance frequency was further calculated from the rate of high organic loading at each disturbance level resulting in values of 0, ^1^/_8_, ^1^/_6_, ^1^/_4_, ^1^/_2_, and 1. Ecosystem function, in the form of process performance parameters at the end of a cycle, was measured weekly in accordance with Standard Methods^56^ where appropriate, and targeted the removal of soluble COD and TKN from the mixed liquor after feeding. Sludge settling capacity was measured via the SVI (mL/g), considering 30 minutes of settling time. Concentrations in the mixed liquor of the bioreactors after feeding (*i.e*, beginning of a new cycle) were regularly 305.8 (±7.4) mg COD/L and 45.6 (±0.8) mg TKN/L, or 594.7 (±18.6) mg COD/L and 46.1 (±0.2) mg TKN/L when double organic loading occurred. A food-to-biomass ratio (F:M) control approach was used as previously described^51^, for which biomass was measured weekly as total suspended solids (TSS) after which sludge wastage was done to target a TSS of 1,500 mg/L. The latter resulted in average solids residence time (SRT) values of 30, 26, 23, 22, 19 and 15 days, for disturbance levels from 0 to 5, respectively. Note that these SRT values are well above the doubling times of relevant bacteria in activated sludge^57^. Sludge samples of 2 mL (m = 184) were collected on the initial day of the study (four samples, taken at random from the inoculum mix) and weekly from each reactor afterwards (180 samples), for DNA extraction as previously described^37^.

### 16S rRNA gene metabarcoding and reads processing

Bacterial 16S rRNA metabarcoding was done in two steps as described in Santillan *et al.*^51^. Primer set 341f/785r targeted the V3-V4 variable regions of the 16S rRNA gene^58^. The libraries were sequenced in-house at SCELSE on an Illumina MiSeq (v.3) with 20% PhiX spike-in, at 300 bp paired-end read-length. Sequenced sample libraries were processed with the *dada2* (v.1.3.3) R-package^33^, allowing inference of ASVs^32^. Illumina adaptors and PCR primers were trimmed prior to quality filtering. Sequences were truncated after 280 and 255 nucleotides for forward and reverse reads, respectively. After truncation, reads with expected error rates higher than 3 and 5 for forward and reverse reads, respectively, were removed. After filtering, error rate learning, ASV inference and denoising, reads were merged with a minimum overlap of 20 bp. Chimeric sequences (0.17% on average) were identified and removed. For a total of 184 samples, an average of 18,086 reads were kept per sample after processing, representing 47% of the average forward input reads. Taxonomy was assigned using the SILVA database (v.132)^59^. Diversity and assembly analyses were carried on both unrarefied and rarefied datasets. To generate the rarefied dataset, samples were rarefied to the lowest number of reads (5,089) in a sample after processing (Fig. S11).

### Bacterial community structure analyses and statistics

All reported P-values for statistical tests in this study were corrected for multiple comparisons using a false discovery rate (FDR) of 5%. Hill diversity indices^27^ were used to quantify taxonomic α-diversity as described elsewhere^17^. Phylogenetic α-diversity was assessed through Faith’s phylogenetic distance^28^ (PD) including its abundance-weighted version (PD_W_). Community structure in terms of taxonomic β-diversity was evaluated through: i) canonical analysis of principal coordinates (CAP) ordination including ellipses of 60% group-average cluster similarity; ii) misclassification error analysis for each disturbance frequency level over the six time points sampled, via the leave-one-out allocation of observations to groups from CAP; and iii) multivariate tests of permutational analysis of variance (PERMANOVA) and permutational analysis of dispersion (PERMDISP); all from Bray-Curtis dissimilarity matrixes at each time point sampled (30 bioreactors, n = 5), constructed from square-root transformed abundance data using PRIMER (v.7)^60^. Phylogenetic β-diversity was assessed via non-metric multidimensional (NMDS) ordination of a weighted Unifrac dissimilarity matrix, constructed from Hellinger transformed abundance data of all 184 samples using the *phyloseq^61^* R-package (v.1.30.0) in R. The *ggplot2* package (v.3.3.2) in R^62^ was used for local polynomial regression fitting via the *loess* function (including 95% confidence intervals) and box plots construction (using Tukey style whiskers). The *ggdist* R-package (v.2.4.1) was used to make the βNTI raincloud plot. Univariate testing through Welch’s analysis of variance (ANOVA) with Games-Howell post-hoc grouping was done using the *rstatix^63^* (v.0.6.0) R-package. Kendall correlations were done using the *ggpubr^64^* package (v.0.4.0) in R. Heat maps for bacterial phyla relative abundances were constructed using the *ampvis2*^65^ package (v.2.6.2) in R.

### Bacterial community assembly analyses and statistics

The effect of underlying assembly mechanisms was assessed using phylogenetic-based null modelling approaches on both α- and β-diversity. First, the nearest taxon index (NTI)^29^ was calculated for each community to assess whether α-diversity was more or less structured than would be expected by random chance. The model uses the mean nearest taxon distance (MNTD)^29^, which quantifies the phylogenetic distance between each ASV in one community, as a measure of the clustering of closely related ASVs. Phylogenetic relatedness of ASVs was characterized by multiple-alignment of ASV sequences using *decipher* (v.2.14.0) R-package^66^. The phylogenetic tree was then constructed and a GTR+G+I maximum likelihood tree was fitted using the *phangorn* (v.2.5.5) R-package^67^. To quantify the degree to which MNTD deviates from a null model expectation, ASVs and abundances were shuffled across the tips of the phylogenetic tree. After shuffling, MNTD was recalculated to obtain a null value, and repeating the shuffling 1,000 times provided a null distribution. Then, NTI was calculated as the difference between the mean of the null distribution and the observed MNTD in units of standard deviation^29^. The closer to zero a NTI value is, the closer to the null expectation (*i.e.*, higher stochasticity) is the phylogenetic dispersion of a given community. Positive NTI values suggest phylogenetic clustering while negative values indicate phylogenetic overdispersion. Second, β-diversity null modelling via the β-nearest taxon index (βNTI) was done to investigate if the phylogenetic turnover across two samples was significantly more or less similar than would be expected by just random chance^30^. The model uses the βmean nearest taxon distance (βMNTD), which quantifies the phylogenetic distance between pairs of ASVs drawn from two distinct communities. To quantify the degree to which βMNTD deviates from a null model expectation, ASVs and abundances were shuffled across the tips of the phylogenetic tree. After shuffling, βMNTD was recalculated to obtain a null value, and repeating the shuffling 1,000 times provided a null distribution. Then, βNTI was calculated as the difference between the mean of the null distribution and the observed βMNTD in units of standard deviation^30^. The closer to zero a βNTI value is, the closer to the null expectation (*i.e.*, higher stochasticity) is the phylogenetic turnover between two communities. By convention, a value of |βNTI| > 2 indicates that the observed turnover is significantly deterministic, while |βNTI| < 2 indicates dominance of stochastic assembly processes^20^. Similarly, here we consider that |NTI| < 2 indicates dominance of stochastic phylogenetic clustering. Both unweighted and abundance-weighted NTI and βNTI values were calculated. These analyses were done using the *metagMisc^68^* (v.0.0.4) and *picante^69^* (v.1.8.2) R-packages. To test for a phylogenetic signal across phylogenetic distances, Mantel correlograms were constructed using the *vegan^70^* (v.2.5.6) R-package, relating between-ASV niche differences to between-ASV phylogenetic distances across a given phylogenetic distance, following the previously described methodology^20,30^. Environmental niches were constructed from bioreactor effluent process data (COD removal, TKN removal and SVI). Phylogenetic distances were quantified for 50 phylogenetic distance bins, while the significance of Pearson correlations was assessed using 1,000 permutations and FDR (5%) correction.

## Acknowledgements

This research was supported by the Singapore National Research Foundation and Ministry of Education under the Research Centre of Excellence Program. We thank JYJ Tan, JQ Teo, A Latiff, SS Thi, AFBM Batcha, CK Aw, ABA Aziz, and JHJ Lim for their assistance with laboratory work. DI Drautz-Moses provided support for the 16S rRNA gene amplicon library preparation and sequencing pipelines employed. We thank J Thompson and VR Regina for helpful suggestions that improved the manuscript.

## Author Contributions

Both authors conceived the idea. ES designed and performed the experiment, as well as data processing and analyses. SW obtained the funding for the study. ES wrote the manuscript draft. Both authors contributed to manuscript editing.

## Data availability

DNA sequencing data are available at NCBI BioProjects with accession number PRJNA723443. See supplementary information for details about the sludge inoculum collection, synthetic feed preparation, and additional figures of diversity and community assembly metrics, correlations, heat maps and data rarefaction. Diversity analyses on rarefied data and all other relevant data to reproduce the results of this study are available as supplementary files in the online version of this manuscript.

