## supplementary information for "Microbiome assembly predictably shapes diversity across a range of disturbance frequencies"

*Sludge inoculum collection and experiment setup*

Sludge inoculum was collected from one of the activated sludge tanks of a water reclamation plant in Singapore, with a Modified Ludzack-Ettinger (MLE) process configuration. Operation parameters were: Q ≈ 200,000 $m^3$/d, T ≈ 30 °C, pH ≈ 6.7, total suspended solids (TSS) ≈ 1,500 mg/L, hydraulic residence time (HRT) ≈ 8-12 h, and solids residence time (SRT) ≈ 5-6 d. Typical influent concentrations were: total Kjeldahl nitrogen (TKN) ≈ 49 mg/L and total chemical oxygen demand (COD) ≈ 320 mg/L. The plant receives a mix of residential, commercial and industrial wastewater as its influent, operating continuously at C:N ≈ 6.5 mg COD/mg TKN and F:M ≈ 0.2-0.3 mg COD/mgTSS/d. It had a removal efficiency of around 80% for N and 90% for COD. On the day the sludge inoculum was collected, the average (± s.d.m. for n = 4) soluble influent concentrations to the secondary treatment were (in units of mg/L): COD = 220.7 ± 2.9, $NH_4^+$-N = 37.4 ± 0.8, TKN = 44.6 ± 0.4, $NO_2^-$-N = 0.00 ± 0.00, $NO_3^-$-N = 0.03 ± 0.00, $PO_4^{3-}$-P = 4.81 ± 0.02. Likewise, the soluble effluent concentrations from the secondary treatment were (in units of mg/L): COD = 34.3 ± 2.1, $NH_4^+$-N = 6.4 ± 0.1, TKN = 8.8 ± 0.1, $NO_2^-$-N = 0.01 ± 0.00, $NO_3^-$-N = 0.03 ± 0.00, $PO_4^{3-}$-P = 0.25 ± 0.04. With these values we estimated an influent COD removal of 0.84 ± 0.01 and an influent TKN removal of 0.80 ± 0.00. Activated sludge was collected in a 20-L container and immediately transported to the lab. The SVI of the inoculum sludge was 108.9 ± 2.2 mL/g, considering 30 min of settling time. The suspension was manually mixed by shaking the closed container thoroughly before transferring half of it to a 10-L vessel that was stirred using a magnetic stir plate to ensure homogeneity. Samples of 25 mL were transferred to thirty 50-mL tubes (Eppendorf), which served as sequencing batch reactors (SBR) in a microcosm setup. About 30 min of settling time was allowed and 12.5 mL of supernatant was removed and replaced with 12.5 mL of synthetic wastewater with or without double organic loading as described below. On the first day a mix of synthetic wastewater with double organic loading was added to reactors for levels 1 to 5, while level 0 reactors received regular synthetic wastewater. All reactors were capped and incubated until the following day in an incubator shaker at 30°C, the prevailing water temperature for wastewater treatment plants in Singapore. After each cycle (24 h) all the tubes were removed from the incubator and allowed to settle for 30 min, after which 12.5 mL of "effluent" supernatant liquid was removed and replaced aseptically with 12.5 mL of fresh synthetic medium, resulting in a 48-h HRT.

*Bioreactor feeding and complex synthetic wastewater preparation*

The composition of the regular synthetic wastewater in the bioreactor feed was adapted from Santillan *et al.*[1]. It contained the following compounds, expressed in mg $L^{-1}$ in mixed liquor in each reactor right after feeding: yeast extract (19.8), soy peptone (18.4), meat peptone (26.3), casein peptone (27.7), sodium acetate anhydrous (119.9), dextrose anhydrous (95.9), urea (37.7), ammonium bicarbonate (33.9), ammonium chloride (63.7), sodium dihydrogen phosphate monohydrate (27.3), sodium phosphate dibasic dihydrate (4.9), calcium chloride dihydrate (50.0), magnesium sulfate heptahydrate (112.5), and sodium bicarbonate (180) which was added to replace the alkalinity consumed during nitrification. The medium also contained 0.25 mL/L of a trace element stock which contained (g/L) citric acid monohydrate (5), EDTA acid disodium salt dihydrate (1.2), hippuric acid (4), sodium molybdate dihydrate (0.24), potassium iodide (0.24), sodium tungstate dihydrate (0.24), boric acid (1), cobalt(II) chloride hexahydrate (0.24), copper(II) sulfate pentahydrate (0.48), manganese(II) chloride tetrahydrate (0.96), nickel(II) chloride hexahydrate (0.24), nitrilotriacetic acid trisodium salt monohydrate (2.88), iron(III) chloride hexahydrate (12), zinc sulfate heptahydrate (1.2).

The above values were used to prepare batches of 2-L bottles of sterile media, a total of six to be used as regular bioreactor feed and three for double organic loading feeding. Average feed concentrations of 305.8 (±7.4) mg COD/L and 45.6 (±0.8) mg TKN/L in the mixed liquor after feeding (*i.e.*, beginning of a new cycle) for reactors. Reactors under double organic loading received double the amount of yeast extract (39.5), soy peptone (36.9), meat peptone (52.7), casein peptone (55.3), sodium acetate anhydrous (239.7), and dextrose anhydrous (191.8), as well as less urea (27.5), ammonium bicarbonate (24.7) and ammonium chloride (46.5) to compensate for the increase in organic TKN. This resulted in average feed concentrations of 594.7 (±18.6) mg COD/L and 46.1 (±0.2) mg TKN/L in the mixed liquor after feeding, when applying double organic loading. Phosphate addition targeted a concentration in mixed liquor of 7.45 (±0.8) mg P/L to obtain a N:P of around 6. The synthetic medium to be used for the whole study was prepared on the same day and filtered through a 0.2-µm pore size filter to avoid contamination. The filtrate was stored at 4 °C for the duration of the study and handled in aseptic conditions.

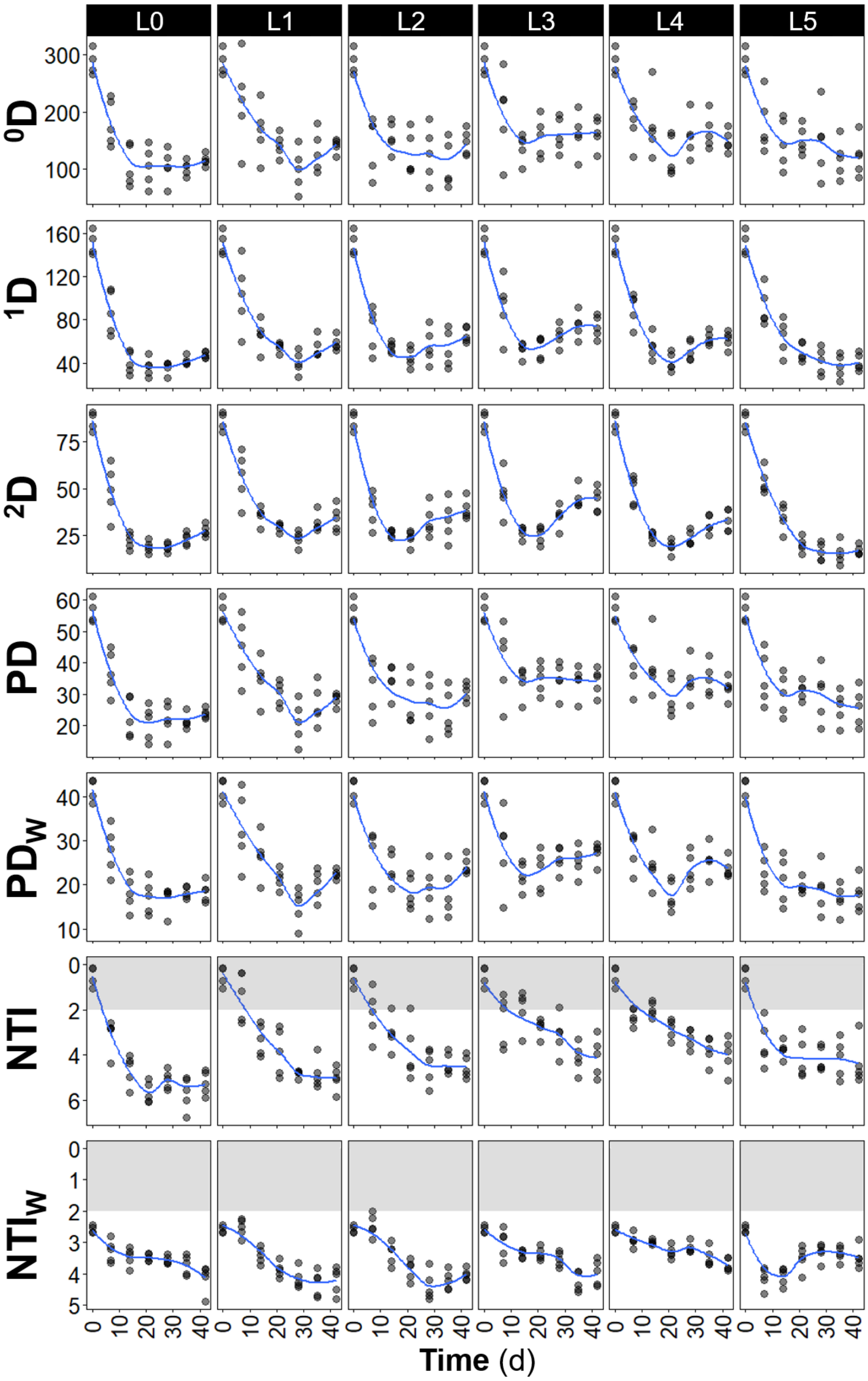


**Fig. S1** – Temporal dynamics of richness ($^0$D), true α-diversity of 1st ($^1$D) and 2nd ($^2$D) order, phylogenetic diversity unweighted (PD) and abundance-weighted ($PD_W$), nearest taxon index unweighted (NTI) and abundance-weighted ($NTI_W$), from bacterial ASV data for each frequency of organic loading disturbance (n = 5, except for day 0 where n = 4). Disturbance frequency levels (L): 0 (undisturbed), 1-4 (intermediately disturbed), 5 (press-disturbed). Blue line represents locally estimated scatterplot smoothing regression (loess). Note the inverted y-axis for both NTI and $NTI_W$, as values closer to zero indicate a higher relative contribution of stochastic assembly. Shaded in grey is the zone of significant stochastic phylogenetic dispersion, $|NTI| < 2$ and $|NTI_W| < 2$.

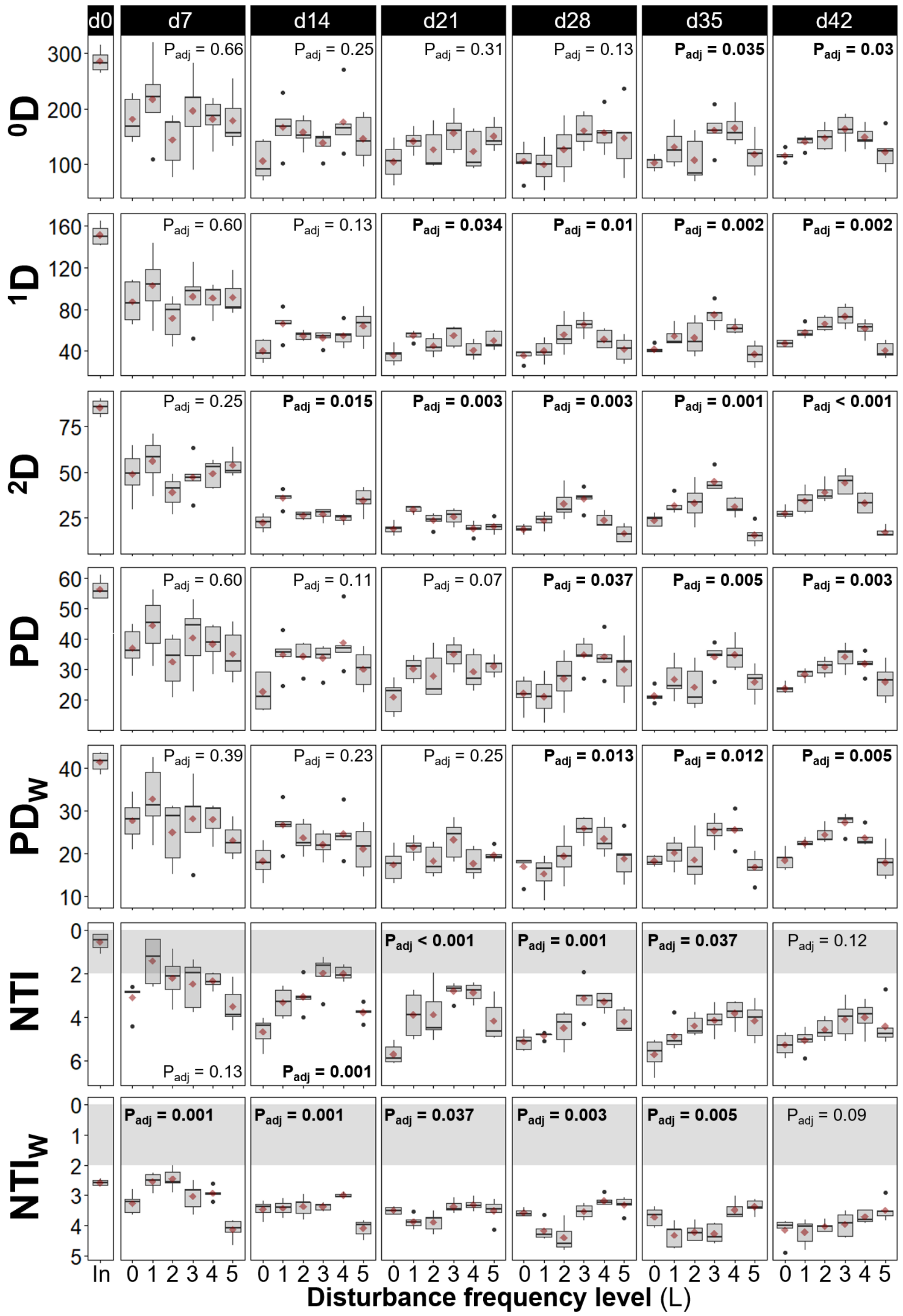


**Fig. S2** – Community structure and assembly assessed via richness ($^0D$), true $\alpha$-diversity of 1st ($^1D$) and 2nd ($^2D$) order, phylogenetic diversity unweighted (PD) and abundance-weighted ($PD_W$), nearest taxon index unweighted (NTI) and abundance-weighted ($NTI_W$), from bacterial ASV data for different frequencies of organic loading disturbance (n = 5). Disturbance frequency levels (L): 0 (undisturbed), 1-4 (intermediately disturbed), 5 (press-disturbed). In: sludge inoculum (day 0, n = 4). Each panel represents a sampling day, red diamonds display mean values. Welch's ANOVA P-values adjusted at 5% FDR shown within panels. Note the inverted y-axis for both NTI and $NTI_W$, as values closer to zero indicate a higher relative contribution of stochastic assembly. Shaded in grey is the zone of significant stochastic phylogenetic dispersion, $|NTI| < 2$ and $|NTI_W| < 2$.

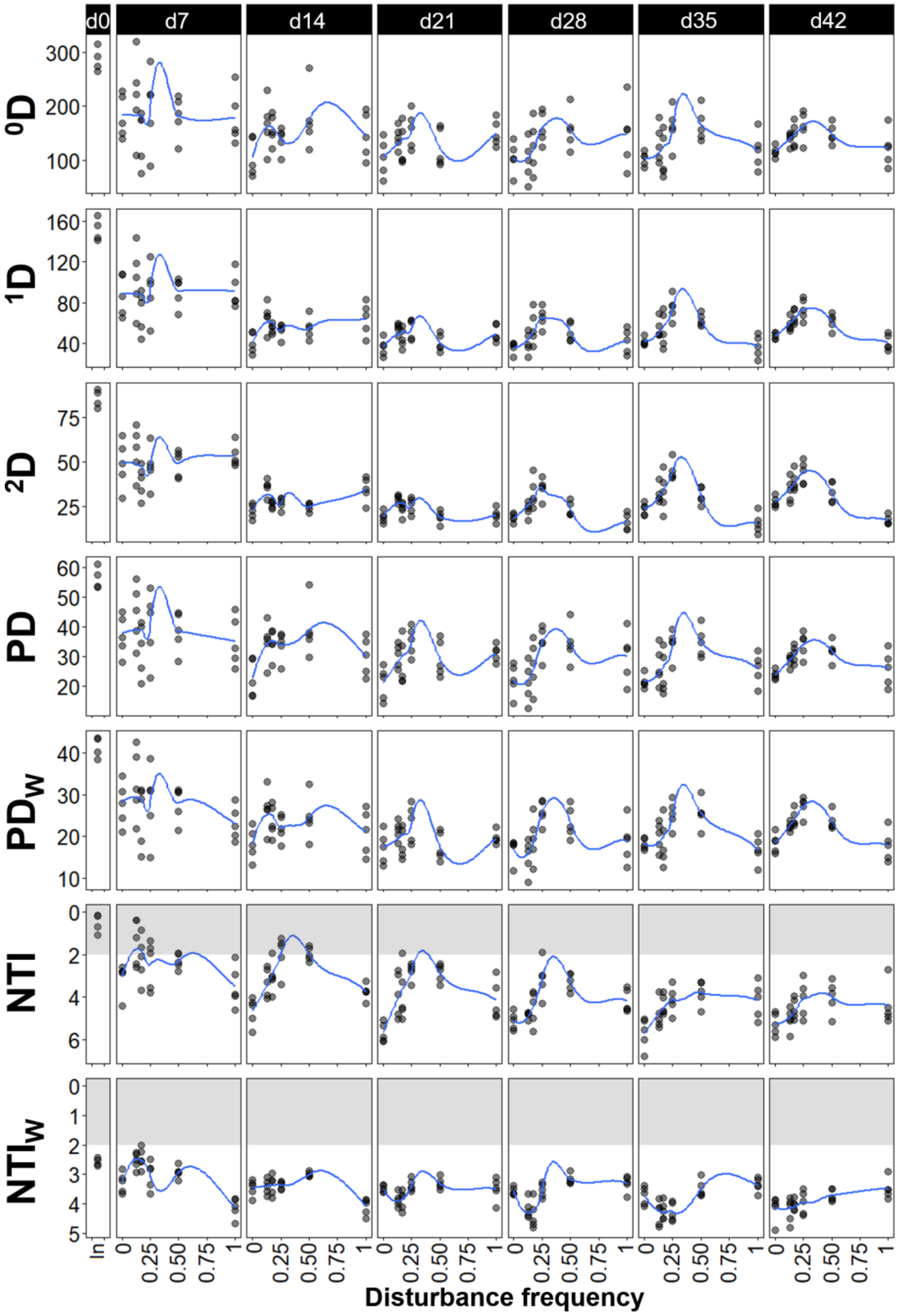


**Fig. S3** – Community structure and assembly assessed via richness ($^0$D), true α-diversity of 1$^{st}$ ($^1$D) and 2$^{nd}$ ($^2$D) order, phylogenetic diversity non-weighed (PD) and abundance-weighed (PD$_W$), nearest taxon index unweighted (NTI) and abundance-weighted (NTI$_W$), from bacterial ASV data for different frequencies of organic loading disturbance (n = 5). Disturbance frequency values were calculated from the frequency of the high organic loading at each disturbance level. In: sludge inoculum (day 0, n = 4). Each panel represents a sampling day. Blue line represents locally estimated scatterplot smoothing regression (loess). Note the inverted y-axis for both NTI and NTI$_W$, as values closer to zero indicate a higher relative contribution of stochastic assembly. Shaded in grey is the zone of significant stochastic phylogenetic dispersion, |NTI| < 2 and |NTI$_W$| < 2.

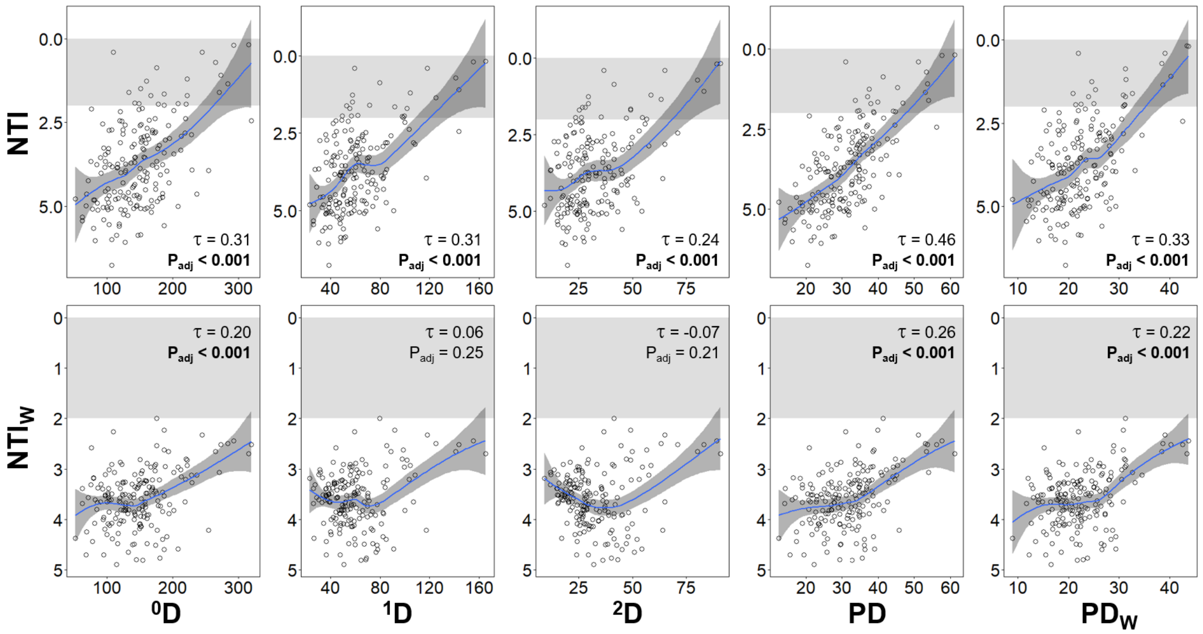


**Fig. S4** – Richness ($^0$D), true α-diversity of 1$^{st}$ ($^1$D) and 2$^{nd}$ ($^2$D) order, unweighted (PD) and abundance-weighted phylogenetic diversity ($PD_W$), correlated against unweighted (NTI, upper panels) and abundance-weighted nearest taxon index ($NTI_W$, lower panels), from bacterial ASV data for all frequency levels and time points evaluated in this study (m = 184). Kendall correlation τ- and P-values adjusted at 5% FDR are indicated within the panels. Blue line represents locally estimated scatterplot smoothing regression (loess) with confidence interval in dark-grey shading. Note the inverted y-axis for both NTI and $NTI_W$, as values closer to zero indicate a higher relative contribution of stochastic assembly. Shaded in grey is the zone of significant stochastic phylogenetic dispersion, |NTI| < 2 and $|NTI_W| < 2$.

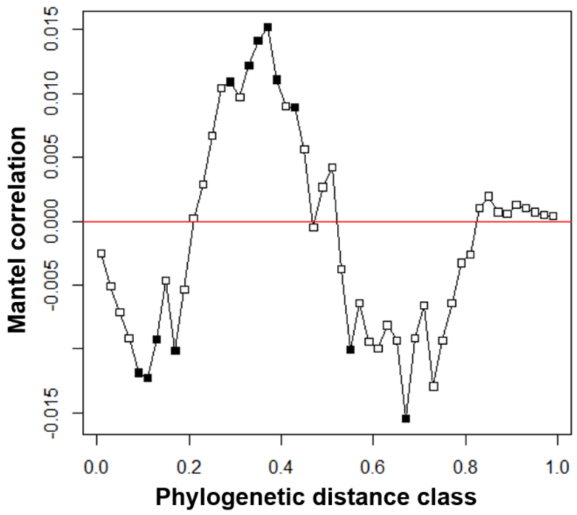


**Fig. S5 -** Phylogenetic Mantel correlogram evaluating phylogenetic signal in the sludge bioreactor communities sampled in this study. The plot relates between-ASV niche differences to between-ASV phylogenetic distances across a given phylogenetic distance. Significant correlations ($P_{adj} < 0.05$; closed symbols) indicate significant phylogenetic signal in ASV ecological niches within the associated phylogenetic distance class. The analysis shows a significant phylogenetic signal but mostly across relatively short phylogenetic distances.

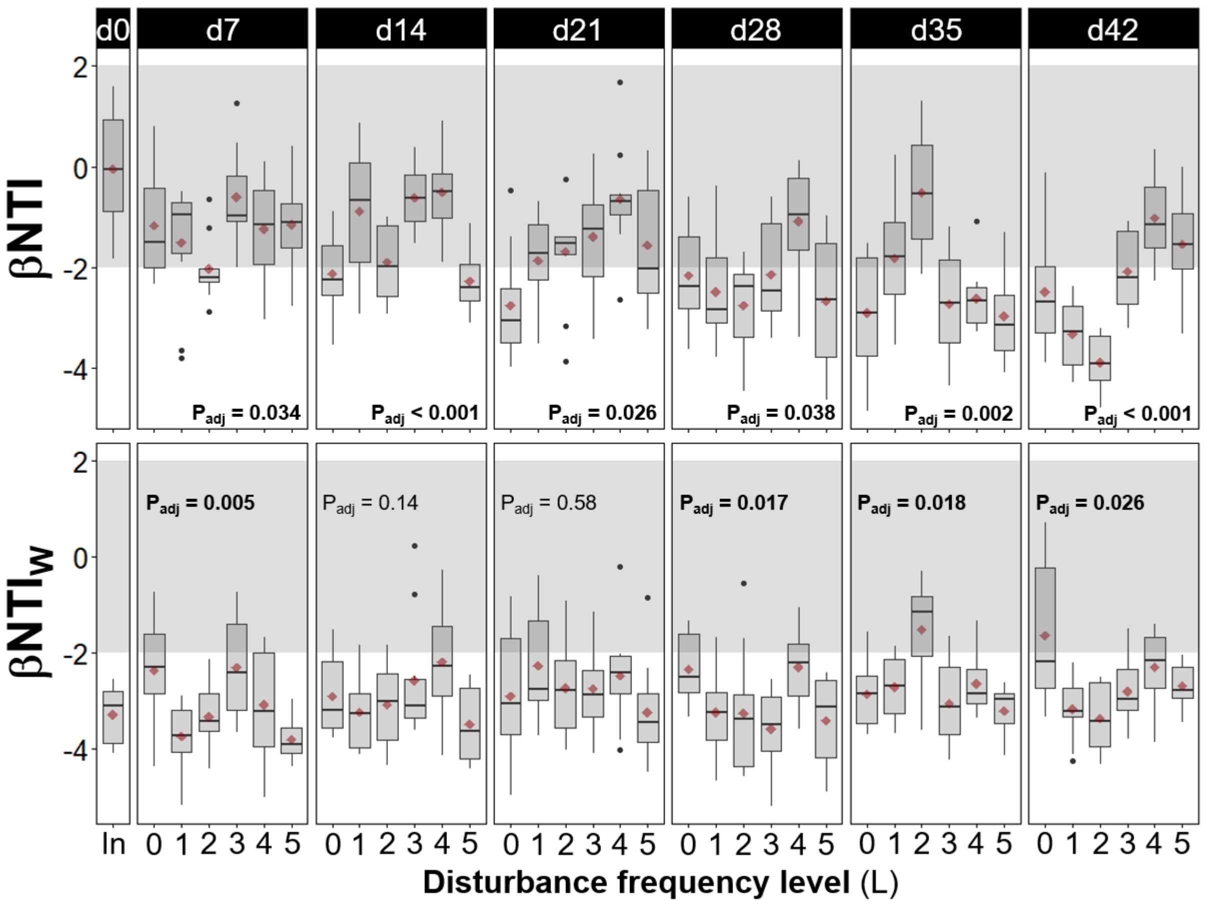


**Fig. S6** – Temporal dynamics of community assembly via β-diversity null modelling of phylogenetic turnover across samples. Within-treatment pairwise values of the β-nearest taxon index, unweighted (βNTI, upper panels) and abundance-weighted ($\beta NTI_W$, lower panels), from bacterial ASV data for different frequencies of organic loading disturbance (n = 10). Disturbance frequency levels (L): 0 (undisturbed), 1-4 (intermediately disturbed), 5 (press-disturbed). In: sludge inoculum (day 0, n = 6). Each panel represents a sampling day, red diamonds display mean values. Welch's ANOVA P-values adjusted at 5% FDR shown within panels. Shaded in grey is the zone where stochastic processes significantly dominate, |βNTI| < 2. βNTI values closer to zero indicate a higher relative contribution of stochastic assembly.

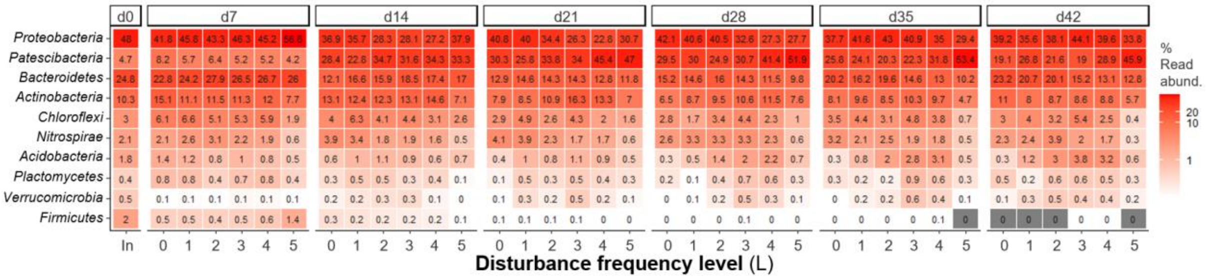


**Fig. S7** – Community structure dynamics for bacterial phyla (rows), assessed through 16S rRNA gene metabarcoding. The 10 most abundant phyla are shown. Columns show the average percentage read abundance among reactors for a given disturbance level across time (n = 5). Disturbance frequency levels (L): 0 (undisturbed), 1-4 (intermediately disturbed), 5 (press-disturbed). In: sludge inoculum (day 0, n = 4).

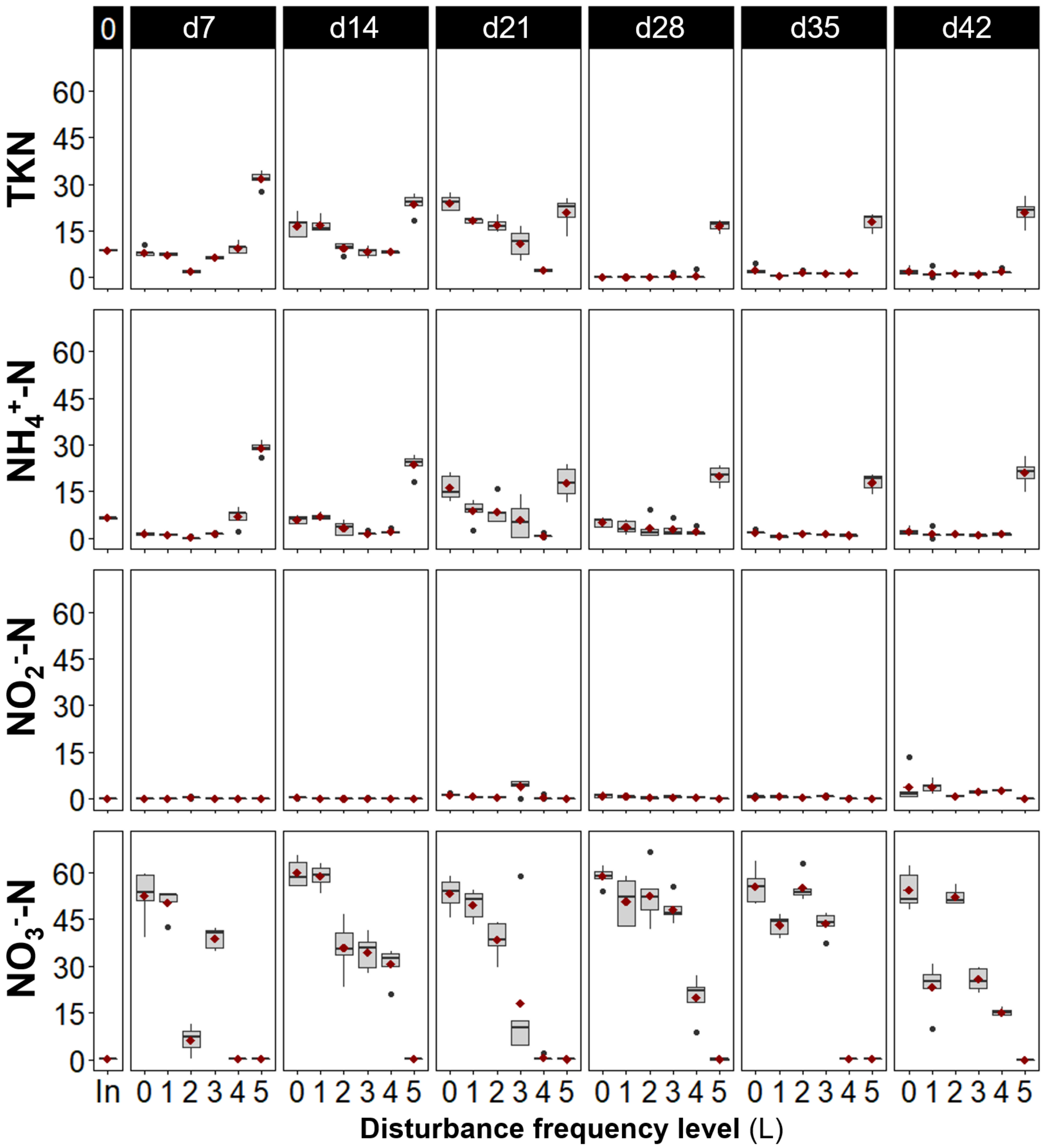


**Fig. S8** – Effluent values of total Kjeldahl nitrogen (TKN), ammonia ($NH_4^+$-N), nitrite ($NO_2^-$-N) and nitrate ($NO_3^-$-N) as nitrogen for different frequencies of organic loading disturbance (n = 5). Disturbance frequency levels (L): 0 (undisturbed), 1-4 (intermediately disturbed), 5 (press-disturbed). In: sludge inoculum (day 0, n = 4). Each panel represents a sampling day, and red diamonds display mean values.

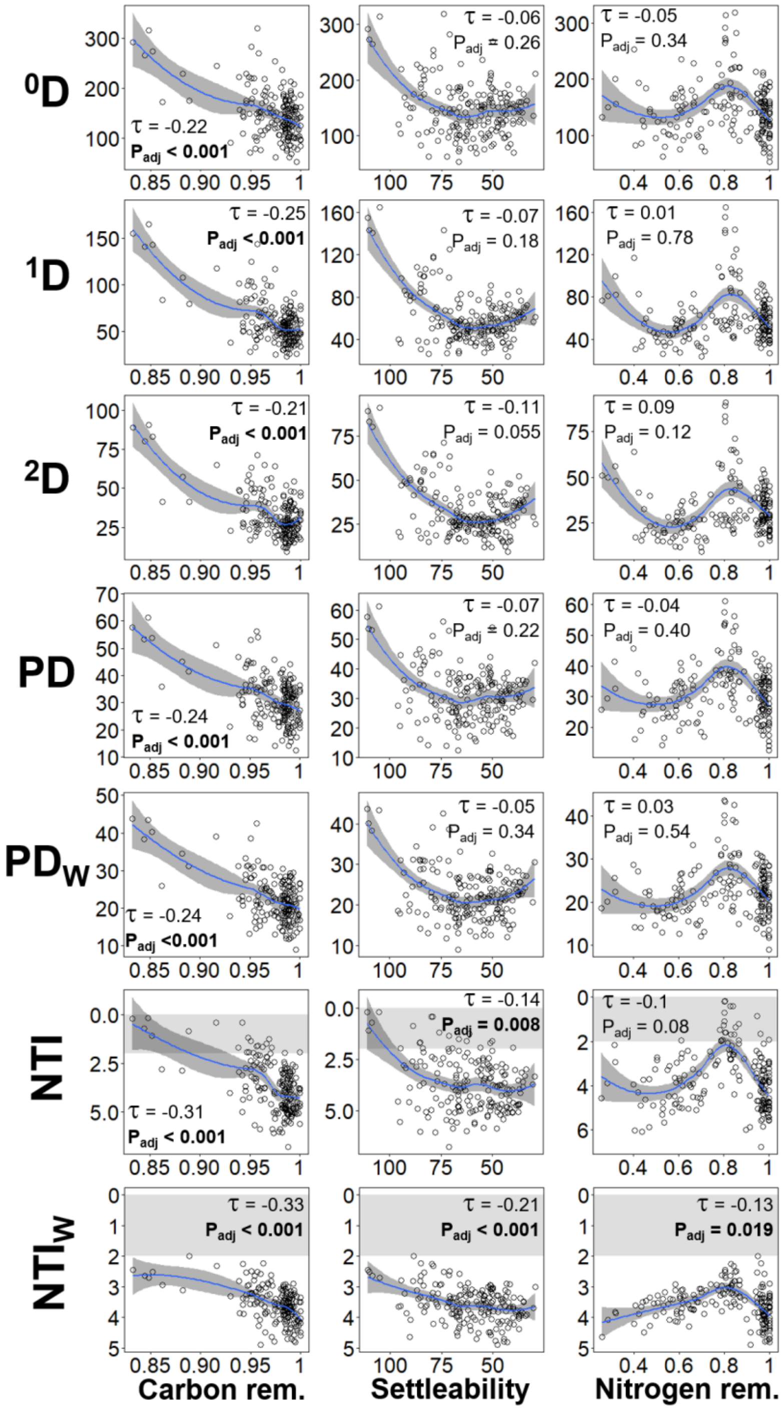


**Fig. S9** – Community function assessed via influent chemical oxygen demand removal (carbon removal, left panels), sludge volume index (sludge settleability, middle panels), and influent total Kjeldahl nitrogen removal (nitrogen removal, right panels), correlated against richness ($^0$D), true α-diversity of 1st ($^1$D) and 2nd ($^2$D) order, unweighted (PD) and abundance-weighted phylogenetic diversity ($PD_W$), unweighted (NTI, upper panels) and abundance-weighted nearest taxon index ($NTI_W$, lower panels), from bacterial ASV data for all frequency levels and time points evaluated in this study (m = 184). Kendall correlation τ- and P-values adjusted at 5% FDR are indicated within the panels. Blue line represents locally estimated scatterplot smoothing regression (loess) with confidence interval in dark-grey shading. Shaded in grey is the zone of significant stochastic phylogenetic dispersion, |NTI| < 2 and $|NTI_W| < 2$. Note the inverted axis for sludge settleability, as it improves with decreasing SVI values, and for both NTI and $NTI_W$, since values closer to zero indicate a higher relative contribution of stochastic assembly.

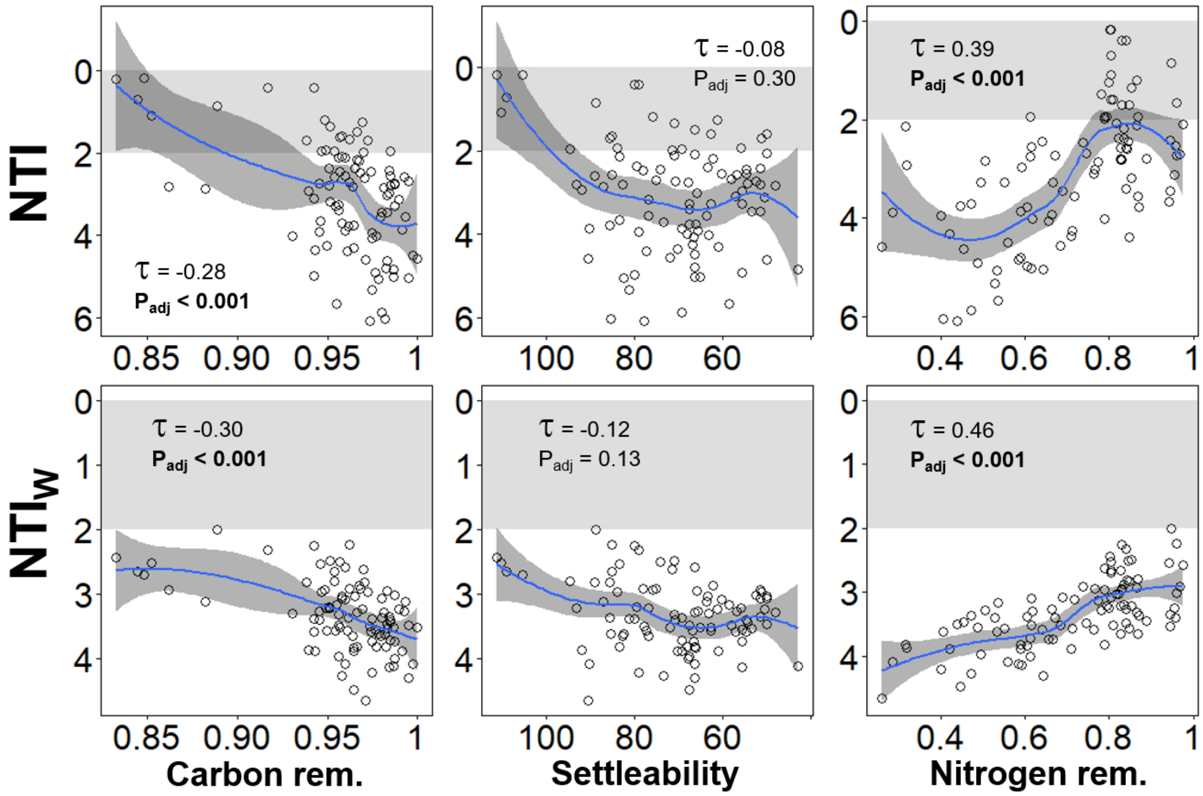


**Fig. S10** – Community function assessed via influent chemical oxygen demand removal (carbon removal, left panels), sludge volume index (sludge settleability, middle panels), and influent total Kjeldahl nitrogen removal (nitrogen removal, right panels), correlated against unweighted (NTI, upper panels) and abundance-weighted nearest taxon index ($NTI_W$, lower panels), from bacterial ASV data at initial stages of succession (d0 to d21, m = 94). Kendall correlation τ- and P-values adjusted at 5% FDR are indicated within the panels. Blue line represents locally estimated scatterplot smoothing regression (loess) with confidence interval in dark-grey shading. Shaded in grey is the zone of significant stochastic phylogenetic dispersion, |NTI| < 2 and $|NTI_W| < 2$. Note the inverted axis for sludge settleability, as it improves with decreasing SVI values, and for both NTI and $NTI_W$, since values closer to zero indicate a higher relative contribution of stochastic assembly.

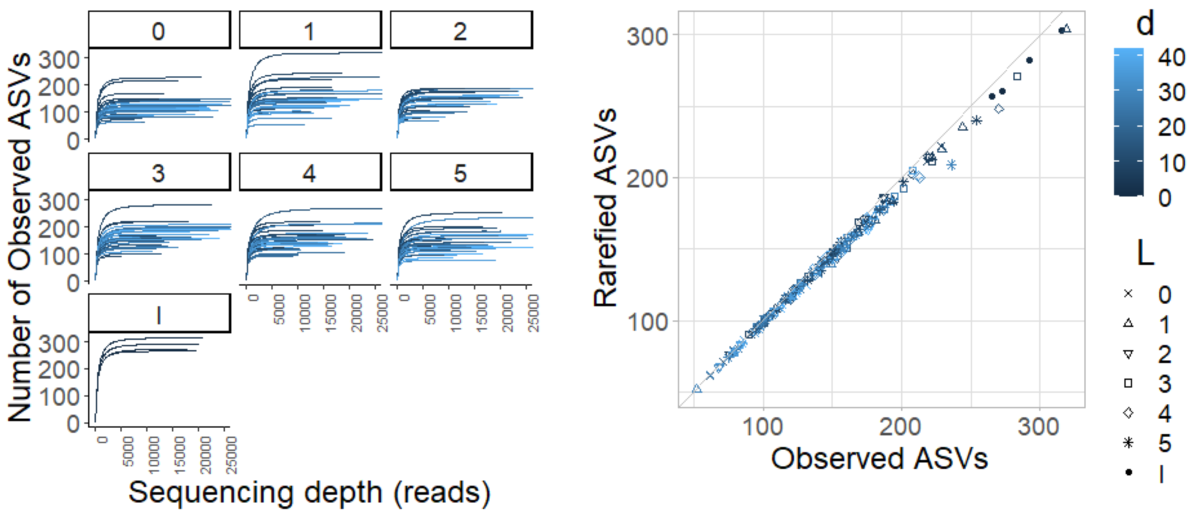


**Fig. S11** – Rarefaction plots for 16S rRNA gene sequencing data. (A) Rarefaction curves for reactors within a given disturbance level across time. (B) Rarefied versus observed number of ASVs. Disturbance frequency levels (L, n = 5): 0 (undisturbed), 1-4 (intermediately disturbed), 5 (press-disturbed). In: Sludge inoculum (day 0, n = 4).

**Table S1.** Multivariate tests on bacterial community structure[Ψ] across disturbance frequency levels.

| **Time** (d) | **No of levels*** | ***n***[‡] | **df**[§] **res** | **PERMANOVA**[†] | | **PERMDISP**[†] | |
|---|---|---|---|---|---|---|---|
| | | | | **F** | **$P_{adj}$**[¶] | **F** | **$P_{adj}$**[¶] |
| 7 | 6 | 5 | 24 | 2.64 | <0.001 | 1.44 | 0.50 |
| 14 | 6 | 5 | 24 | 5.19 | <0.001 | 1.88 | 0.45 |
| 21 | 6 | 5 | 24 | 8.13 | <0.001 | 2.66 | 0.13 |
| 28 | 6 | 5 | 24 | 9.75 | <0.001 | 2.70 | 0.13 |
| 35 | 6 | 5 | 24 | 9.23 | <0.001 | 0.67 | 0.92 |
| 42 | 6 | 5 | 24 | 11.9 | <0.001 | 0.52 | 0.90 |

[Ψ]The Bray-Curtis dissimilarity metric was used on square-root transformed ASV relative abundance data.

* Factor levels (L): 0 (undisturbed), 1-4 (intermediately disturbed), 5 (press-disturbed).

[†] Number of permutations used was 9,999

[‡] Number of replicates per level

[§] Degrees of freedom of the residual

[¶] P-values after correction for multiple comparisons at 5% FDR, via the Benjamini-Hochberg's method.
