## supplementary file for "Microbiome assembly predictably shapes diversity across a range of disturbance frequencies"

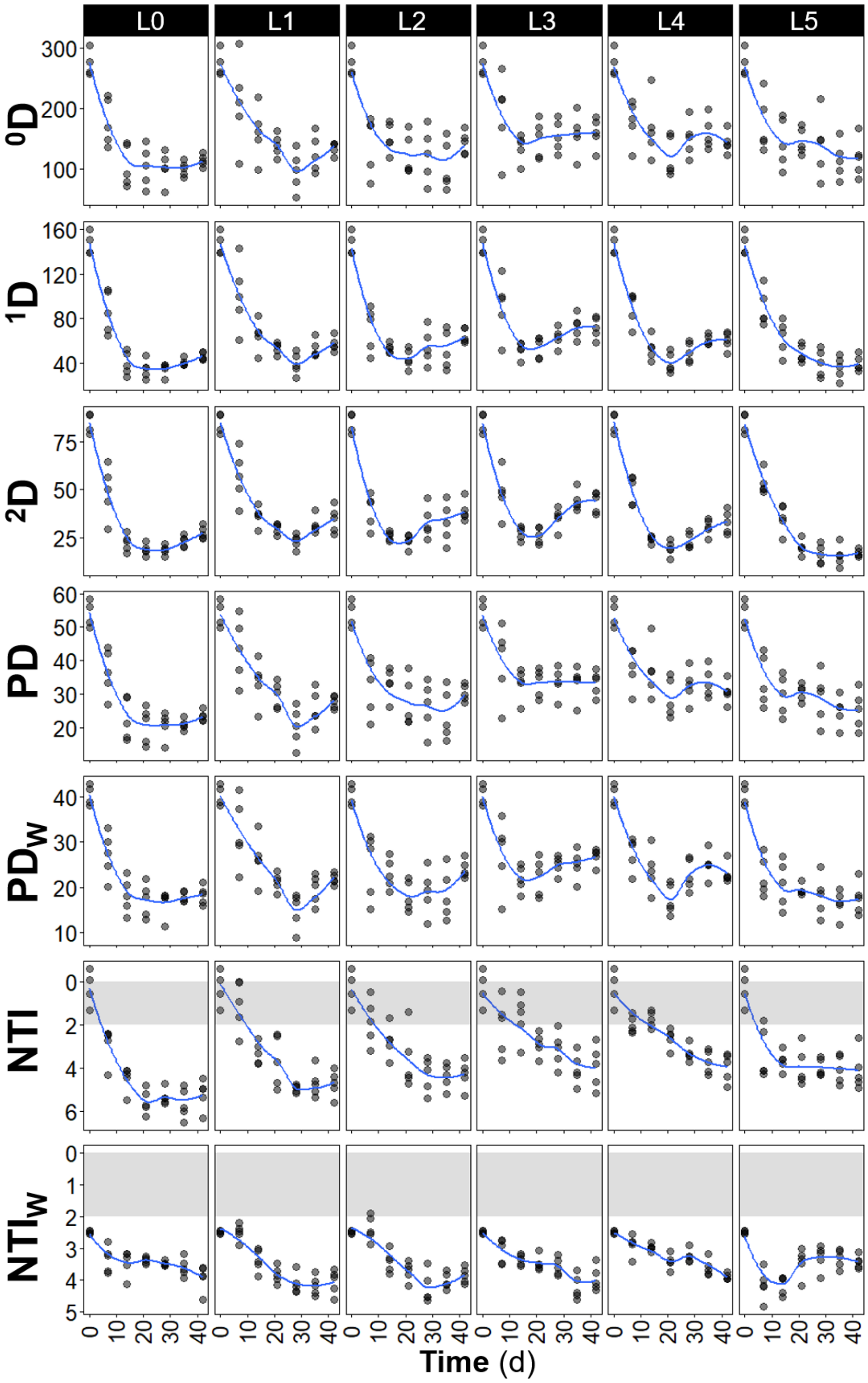


**Fig. SF1** – Temporal dynamics of richness ($^0$D), true α-diversity of 1$^{st}$ ($^1$D) and 2$^{nd}$ ($^2$D) order, phylogenetic diversity unweighted (PD) and abundance-weighted ($PD_W$), nearest taxon index unweighted (NTI) and abundance-weighted ($NTI_W$), from **rarefied** bacterial ASV data for each frequency of organic loading disturbance (n = 5, except for day 0 where n = 4). Disturbance frequency levels (L): 0 (undisturbed), 1-4 (intermediately disturbed), 5 (press-disturbed). Blue line represents locally estimated scatterplot smoothing regression (loess). Note the inverted y-axis for both NTI and $NTI_W$, as values closer to zero indicate a higher relative contribution of stochastic assembly. Shaded in grey is the zone of significant stochastic phylogenetic dispersion, |NTI| < 2 and |$NTI_W$| < 2.

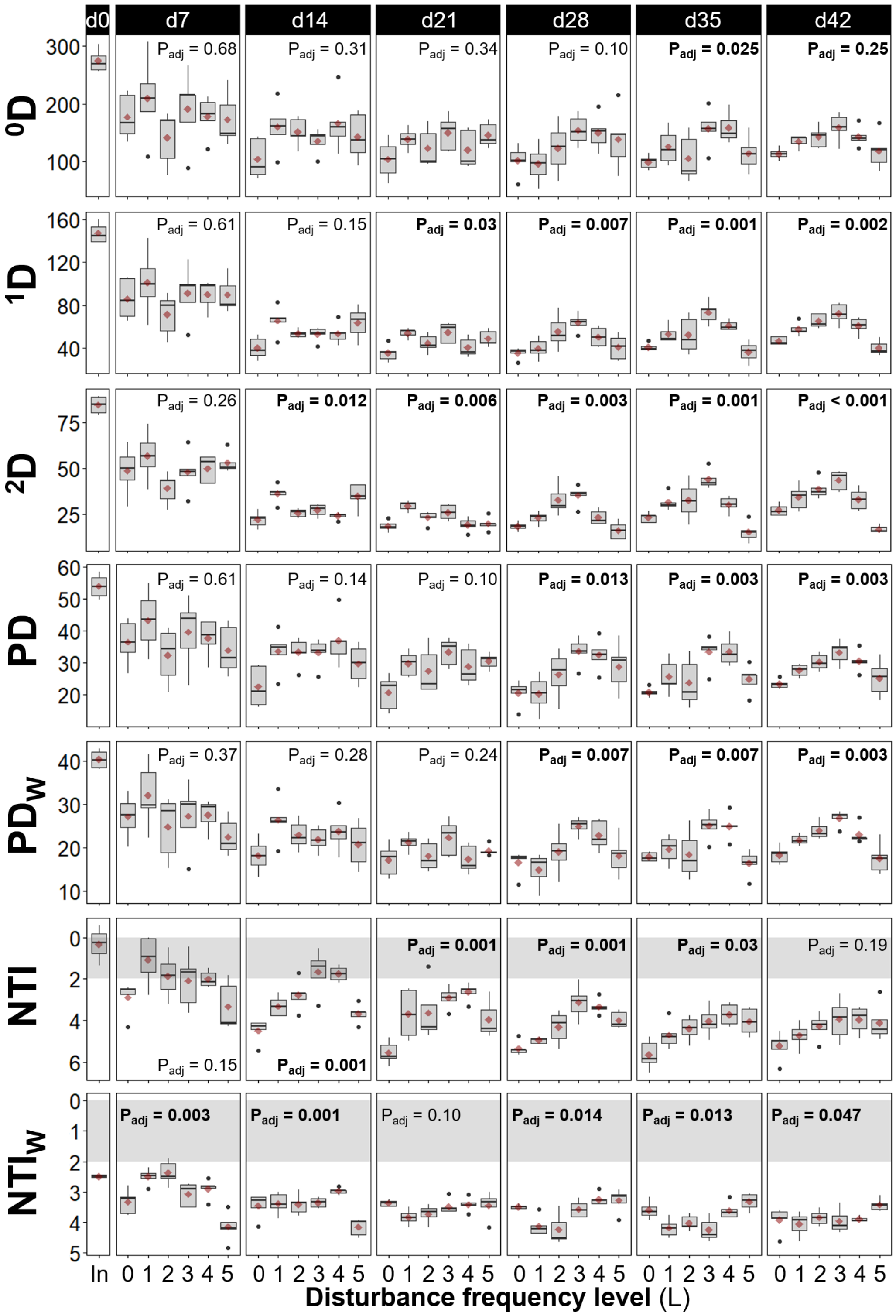


**Fig. SF2** – Community structure and assembly assessed via richness ($^0$D), true α-diversity of 1st ($^1$D) and 2nd ($^2$D) order, phylogenetic diversity unweighted (PD) and abundance-weighted ($PD_W$), nearest taxon index unweighted (NTI) and abundance-weighted ($NTI_W$), from **rarefied** bacterial ASV data for different frequencies of organic loading disturbance (n = 5). Disturbance frequency levels (L): 0 (undisturbed), 1-4 (intermediately disturbed), 5 (press-disturbed). In: sludge inoculum (day 0, n = 4). Each panel represents a sampling day, red diamonds display mean values. Welch's ANOVA P-values adjusted at 5% FDR shown within panels. Note the inverted y-axis for both NTI and $NTI_W$, as values closer to zero indicate a higher relative contribution of stochastic assembly. Shaded in grey is the zone of significant stochastic phylogenetic dispersion, $|NTI| < 2$ and $|NTI_W| < 2$.

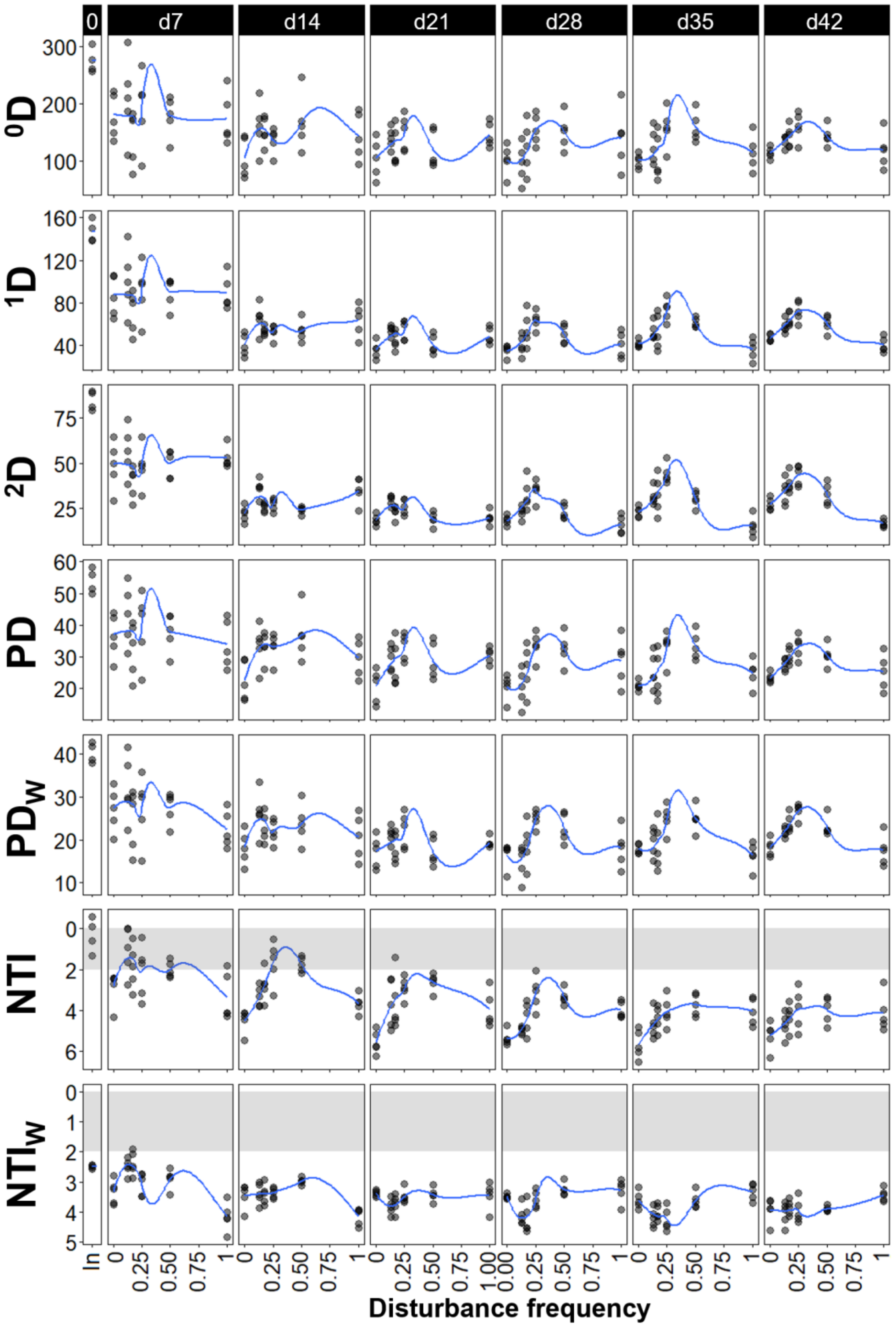


**Fig. SF3** – Community structure and assembly assessed via richness (${}^{0}$D), true α-diversity of 1st (${}^{1}$D) and 2nd (${}^{2}$D) order, phylogenetic diversity non-weighed (PD) and abundance-weighed ($PD_W$), nearest taxon index unweighted (NTI) and abundance-weighted ($NTI_W$), from **rarefied** bacterial ASV data for different frequencies of organic loading disturbance (n = 5). Disturbance frequency values were calculated from the frequency of the high organic loading at each disturbance level. In: sludge inoculum (day 0, n = 4). Each panel represents a sampling day. Blue line represents locally estimated scatterplot smoothing regression (loess). Note the inverted y-axis for both NTI and $NTI_W$, as values closer to zero indicate a higher relative contribution of stochastic assembly. Shaded in grey is the zone of significant stochastic phylogenetic dispersion, $|NTI| < 2$ and $|NTI_W| < 2$.

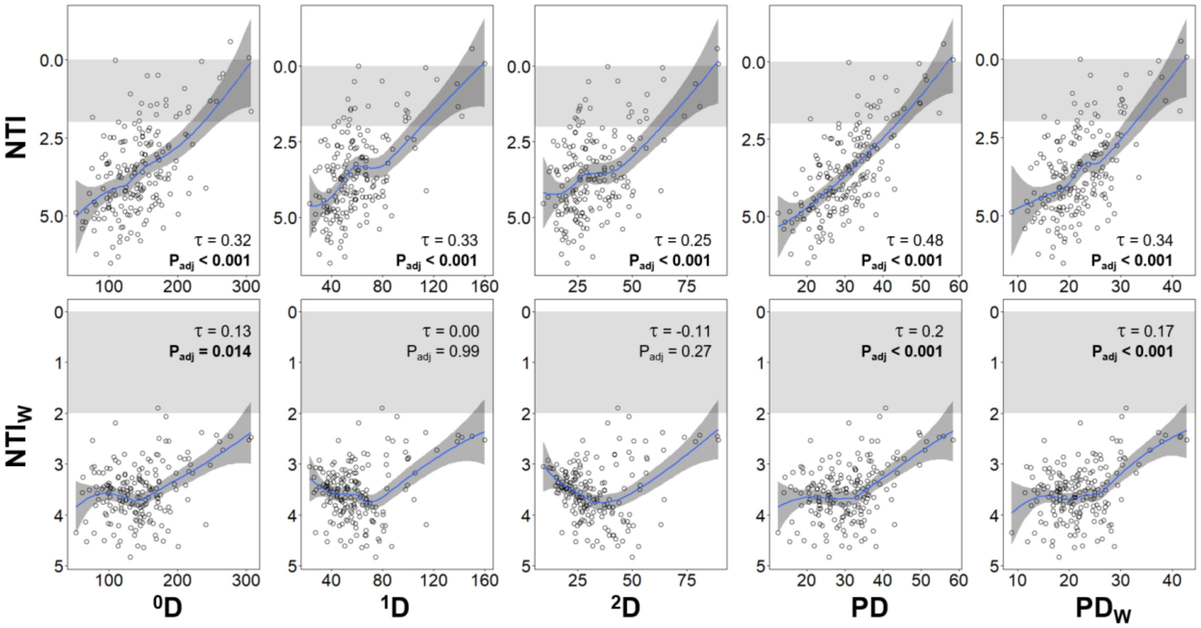


**Fig. SF4** – Richness ($^0$D), true α-diversity of 1$^{st}$ ($^1$D) and 2$^{nd}$ ($^2$D) order, unweighted (PD) and abundance-weighted phylogenetic diversity ($PD_W$), correlated against unweighted (NTI, upper panels) and abundance-weighted nearest taxon index ($NTI_W$, lower panels), from **rarefied** bacterial ASV data for all frequency levels and time points evaluated in this study (m = 184). Kendall correlation τ- and P-values adjusted at 5% FDR are indicated within the panels. Blue line represents locally estimated scatterplot smoothing regression (loess) with confidence interval in dark-grey shading. Note the inverted y-axis for both NTI and $NTI_W$, as values closer to zero indicate a higher relative contribution of stochastic assembly. Shaded in grey is the zone of significant stochastic phylogenetic dispersion, $|NTI| < 2$ and $|NTI_W| < 2$.

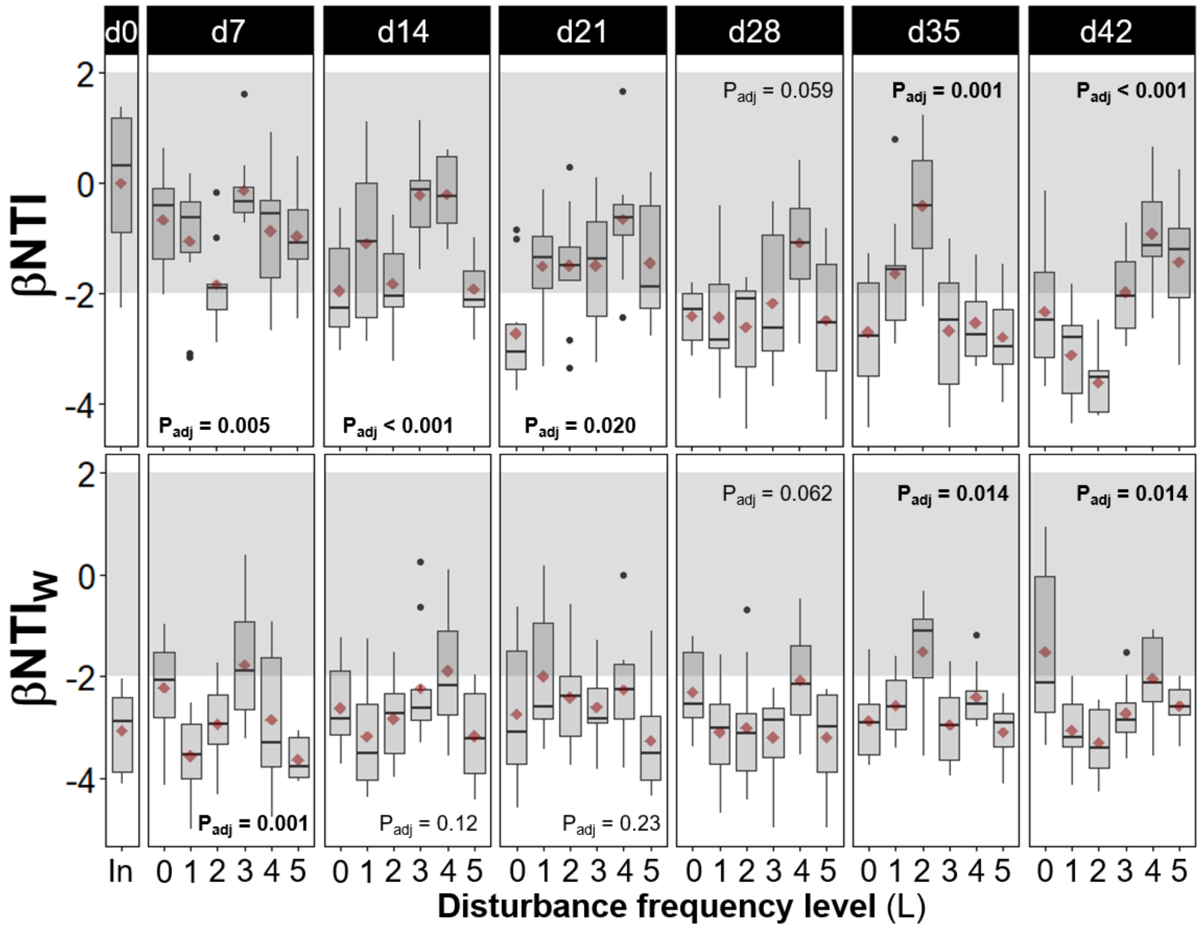


**Fig. SF6** – Temporal dynamics of community assembly via β-diversity null modelling of phylogenetic turnover across samples. Within-treatment pairwise values of the β-nearest taxon index, unweighted (βNTI, upper panels) and abundance-weighted ($βNTI_W$, lower panels), from **rarefied** bacterial ASV data for different frequencies of organic loading disturbance (n = 10). Disturbance frequency levels (L): 0 (undisturbed), 1-4 (intermediately disturbed), 5 (press-disturbed). In: sludge inoculum (day 0, n = 6). Each panel represents a sampling day, red diamonds display mean values. Welch's ANOVA P-values adjusted at 5% FDR shown within panels. Shaded in grey is the zone where stochastic processes significantly dominate, |βNTI| < 2. βNTI values closer to zero indicate a higher relative contribution of stochastic assembly.

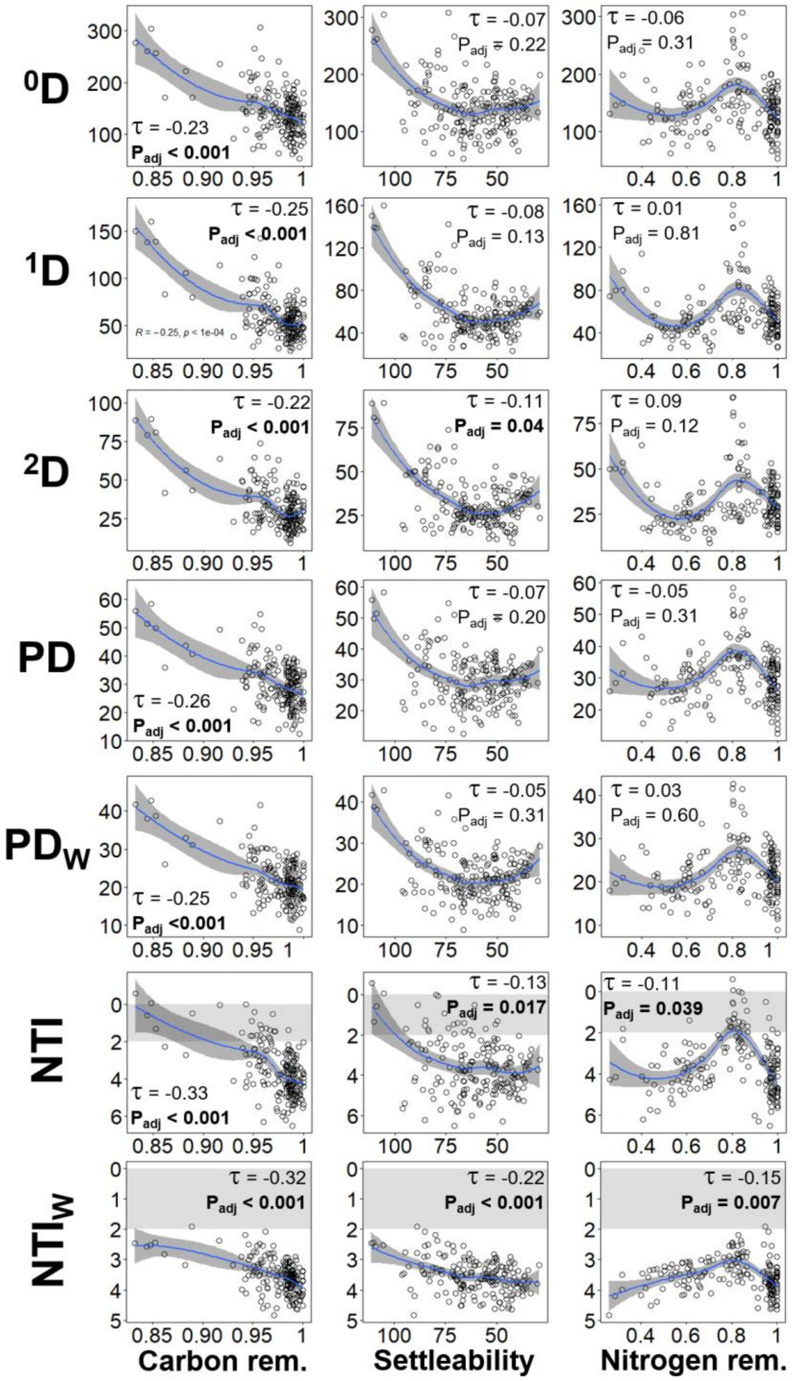


**Fig. SF9** – Community function assessed via influent chemical oxygen demand removal (carbon removal, left panels), sludge volume index (sludge settleability, middle panels), and influent total Kjeldahl nitrogen removal (nitrogen removal, right panels), correlated against richness ($^0D$), true $\alpha$-diversity of 1st ($^1D$) and 2nd ($^2D$) order, unweighted (PD) and abundance-weighted phylogenetic diversity ($PD_W$), unweighted (NTI, upper panels) and abundance-weighted nearest taxon index ($NTI_W$, lower panels), from **rarefied** bacterial ASV data for all frequency levels and time points evaluated in this study (m = 184). Kendall correlation $\tau$- and P-values adjusted at 5% FDR are indicated within the panels. Blue line represents locally estimated scatterplot smoothing regression (loess) with confidence interval in dark-grey shading. Shaded in grey is the zone of significant stochastic phylogenetic dispersion, $|NTI| < 2$ and $|NTI_W| < 2$. Note the inverted axis for sludge settleability, as it improves with decreasing SVI values, and for both NTI and $NTI_W$, since values closer to zero indicate a higher relative contribution of stochastic assembly.

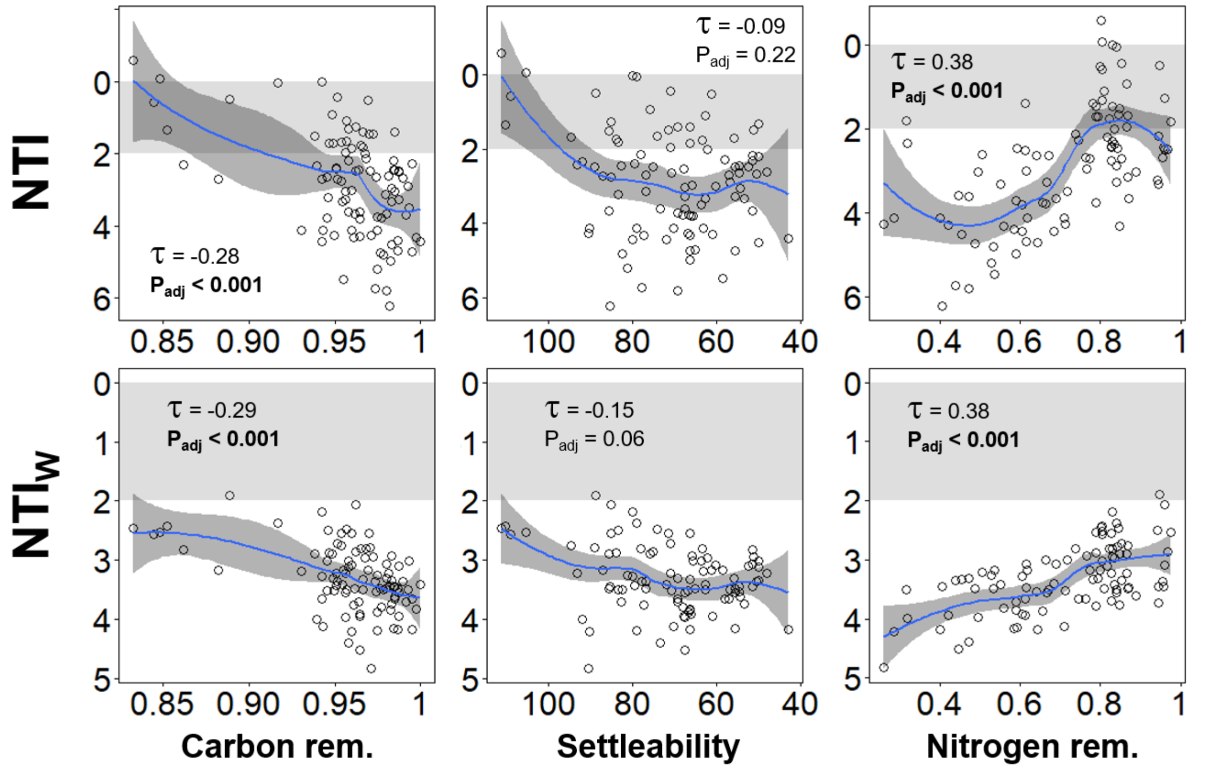


**Fig. SF10** – Community function assessed via influent chemical oxygen demand removal (carbon removal, left panels), sludge volume index (sludge settleability, middle panels), and influent total Kjeldahl nitrogen removal (nitrogen removal, right panels), correlated against unweighted (NTI, upper panels) and abundance-weighted nearest taxon index ($NTI_W$, lower panels), from **rarefied** bacterial ASV data at initial stages of succession (d0 to d21, m = 94). Kendall correlation τ- and P-values adjusted at 5% FDR are indicated within the panels. Blue line represents locally estimated scatterplot smoothing regression (loess) with confidence interval in dark-grey shading. Shaded in grey is the zone of significant stochastic phylogenetic dispersion, $|NTI| < 2$ and $|NTI_W| < 2$. Note the inverted axis for sludge settleability, as it improves with decreasing SVI values, and for both NTI and $NTI_W$, since values closer to zero indicate a higher relative contribution of stochastic assembly.
